# Metal Ion Sensing by Tetraloop-Like RNA Fragment: Role of Compact Intermediates with Non-Native Metal Ion-RNA Inner Shell Contacts

**DOI:** 10.1101/2024.09.25.615097

**Authors:** Sk Habibullah, Lipika Baidya, Sunil Kumar, Govardhan Reddy

**Affiliations:** Solid State and Structural Chemistry Unit, Indian Institute of Science, Bengaluru 560012, Karnataka, India

## Abstract

Divalent metal ions influence the folding and function of RNA in the cells. The mechanism of how RNA structural elements in riboswitches sense specific metal ions is unclear. RNA interacts with ions through two distinct binding modes: direct interaction between the ion and RNA (inner-shell (IS) coordination) and indirect interaction between the ion and RNA mediated through water molecules (outer-shell (OS) coordination). To understand how RNA senses metal ions such as Mg^2+^ and Ca^2+^, we studied the folding of a small RNA segment from the Mg^2+^ sensing M-Box riboswitch using computer simulations. This RNA segment has the characteristics of a GNRA tetraloop motif and interestingly requires binding of a single Mg^2+^ ion. The folding free energy surface of this simple tetraloop system is multidimensional, with a population of multiple intermediates where the tetraloop and cation interact through IS and OS coordination. The partially folded compact tetraloop intermediates form multiple non-native IS contacts with the metal ion. Thermal fluctuations should break these strong non-native IS contacts so that the tetraloop can fold to the native state, resulting in higher folding free energy barriers. Ca^2+^ undergoes rapid OS to IS transitions and vice-versa due to its lower charge density than Mg^2+^. However, the ability of Ca^2+^ to stabilize the native tetraloop state is weaker as it could not hold the loop-closing nucleotides together due to its weaker interactions with the nucleotides. These insights are critical to understanding the specific ion sensing mechanisms in riboswitches, and the predictions are amenable for verification by NMR experiments.

## Introduction

Metal ions are essential for the folding of ribonucleic acid (RNA) into its native specific three-dimensional structure, which is critical for its diverse functions in protein synthesis, ^1^ catalytic activity, ^2^ gene regulation,^3,4^ and gene editing. ^5,6^ Since RNA is a polyelectrolyte with a negatively charged phosphate backbone, it requires counterions to offset the electro-static repulsion among phosphate groups for the close packing of backbone.^7–12^ Intracellular monovalent (Na^+^, K^+^) and divalent cations (Mg^2+^, Ca^2+^, Co^2+^, Ni^2+^, Mn^2+^) act as the counterions that interact with the RNA backbone.^13–19^ However, unlike monovalent ions, the specific non-uniform binding of divalent cations is crucial for RNA structural stability and functionality. ^20–22^ Cation binding to RNA occurs through two distinct modes: 1) inner-shell (IS) contact, where the RNA atom is within the first solvation shell of the ion, and 2) outer-shell (OS) contact, where a water molecule mediates contact between the ion and an RNA atom, and the RNA atom is in the second solvation shell of the ion (Fig. 1A).^23–27^

**Figure 1.**
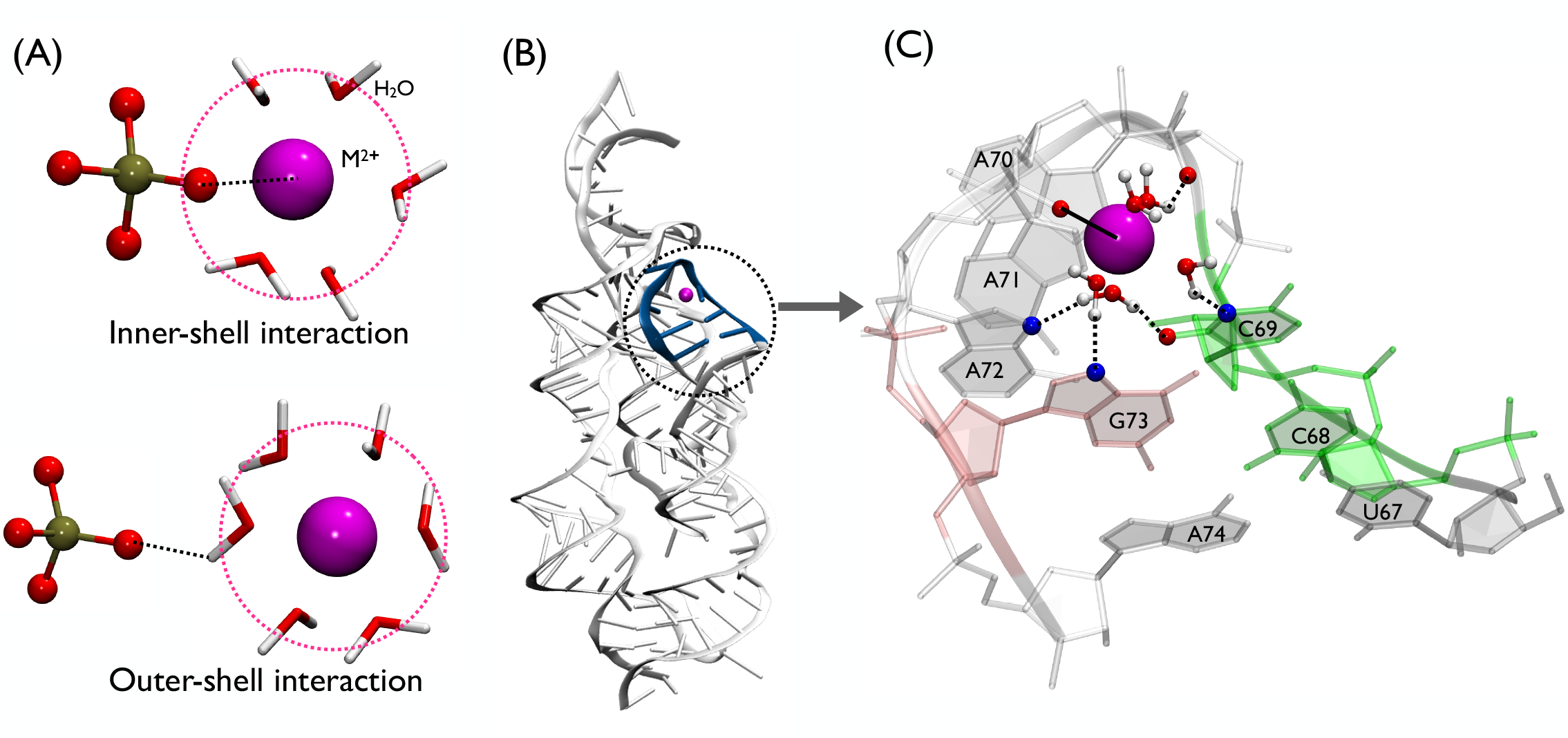
(A) IS interaction is formed through direct contact between the metal ion (M^2+^) and the electronegative atom of RNA. OS interaction occurs through the water-mediated hydrogen bond between the RNA electronegative atom and the water molecule present in the first solvation shell of the metal ion. Tan and red color spheres represent the phosphorus and oxygen atoms in the phosphate group of RNA. Water molecules are shown in the Licorice representation, and the metal ion is shown as a magenta sphere. (B) Crystal structure of the M-box riboswitch (PDB ID -2QBZ)^16^ (white) with the L4 tetraloop shown in blue and is encircled with a black dotted line. (C) Eight nucleotides long L4 tetraloop. Mg^2+^ is shown as a magenta sphere. The bound Mg^2+^ interacts with the phosphate oxygen (O2P) atom of nucleotide A72 through IS contact shown as a black solid line (see Fig. S1A). Atoms involved in OS contact shown as black dotted lines with Mg^2+^ are phosphate oxygen O2P of A71, base oxygen O2 and base nitrogen N3 of C69, base nitrogen N7 of G73 and base nitrogen N7 of A72.

There have been extensive research efforts to understand the role of IS and OS modes of metal ion binding in RNA folding and its functions. ^28–33^ X-ray crystallography and NMR experiments showed that IS/OS-coordinated ions in RNA structures are involved in specific tertiary contact formation through H-bonding and stacking interaction. ^34–37^ In riboswitches, in addition to facilitating folding, Mg^2+^ through IS and OS interaction modes plays a critical role in aiding the binding of negatively charged cognate ligands such as TPP^21,38^ and F^*−*^.^39,40^ Mg^2+^ ions through OS interaction modes are shown to play a role in widening the ligand binding pocket in the SAM riboswitch, aiding in ligand binding.^41^ Further, one of the reasons for the remarkable ability of metal ion-sensing riboswitches to selectively recognize and bind with different metal ions, such as Mg^2+^, Co^2+^, Mn^2+^, and Ni^2+^ is due to the subtle differences in the ability of various ions to form IS and OS interactions in their binding pockets. ^16–19^ Additionally, specific metal ion binding by ribozymes is critical for their catalytic and splicing activity.^42^ Experiments show that substituting Mg^2+^ with Ca^2+^ ions can lead to a significant reduction in ribozyme’s catalytic activity,^43,44^ which is due to the differences in the metal ion binding affinities and water solvation. ^45^ Therefore, it is essential to understand the various factors that contribute to metal ion sensing by RNA.

Earlier studies have emphasized that the driving force for specific metal ion binding to large RNA complexes is the overall gain in structural stabilization due to the tertiary contact formation. ^7,21,22^ However, the mechanism by which a cation binds to a specific RNA pocket, the barriers the ion has to overcome in the binding process, and the reasons for the ion specificity remain elusive. To address this problem, we utilized an RNA segment with the structure of a simple RNA tetraloop devoid of tertiary structural elements and required the binding of a single Mg^2+^ ion for folding. The simplicity of the structure eliminates the complexity originating from the folding of large RNA structures with multiple secondary and tertiary structural elements and their binding to multiple ions. These advantages allow us to probe the binding mechanism of a single metal ion and its role in stabilizing the tetraloop-like structure.

RNA tetraloops are small secondary structural elements present in RNA hairpin motifs formed by four nucleotides, GNRA or UNCG, where N represents any nucleotide, and R represents a purine residue (A or G). In addition to these canonical sequences, multiple other sequences are found to fold to tetraloop-like structures. ^46^ Tetraloops play a significant role in RNA folding, interactions with other biomolecules and tertiary contact formation. ^47–54^ Due to their smaller size and importance in RNA folding and function, they are widely studied using computer simulations. ^20,31,55–61^ Previous simulations on GAGA and UUCG tetraloops revealed the drawbacks in the RNA force fields^55,56,60,61^ and contributed to their improvement.^62,63^

In this work, we studied the folding of a special tetraloop taken from the L4 loop of the M-box riboswitch (PDB ID: 2QBZ)^16^ (Fig. 1B,C). Although the L4 tetraloop sequence (CAAA) is different from other typical RNA tetraloops (GNRA or UNCG), it still adopts the characteristic shape of a GNRA tetraloop. In contrast to the GNRA tetraloop, an Mg^2+^ is bound approximately at the center of the L4 tetraloop. The bound Mg^2+^ ion interacts with the phosphate oxygen atoms of A72 nucleotide in the L4 tetraloop through IS coordination (native ion-tetraloop IS contact) (Fig. 1C and S1A) and the bound Mg^2+^ ion also facilitates the interaction between the loop-closing bases (C69 and A72) through water-mediated hydrogen bonds stabilizing the structure similar to the conventional GNRA tetraloop (Fig. S1A,B). The presence of a single IS-coordinated Mg^2+^ in the folded structure makes the L4 tetraloop an ideal model system to understand the driving forces for metal ion sensing by a small RNA secondary structural element and its folding.

The previous simulation study^20^ on the L4 tetraloop using long unbiased molecular dynamics (MD) simulations showed that monovalent (Na^+^ or K^+^) or divalent (Mg^2+^ or Ca^2+^) metal ions are required for this tetraloop to fold. This study further showed that Mg^2+^ better stabilized the folded tetraloop structure due to its rigid water solvation shell consisting of six water molecules arranged in an octahedral arrangement. However, the folded state was observed to be unstable, and its average lifetime was only *≈* 8.5 ns in the presence of 65 mM Mg^2+^. In these simulations, the OS to IS transition of Mg^2+^ ion and its IS coordination with the phosphate oxygen atom of nucleotide A72 was not observed, which probably accounts for the low stability of the L4 tetraloop folded states observed in these simulations. The reason for not observing the OS to IS transition for Mg^2+^ in these unbiased all-atom simulations could be that the barrier for observing this transition is high.

In this work, we performed all-atom MD simulations of the L4 tetraloop in the presence of divalent cations Mg^2+^ or Ca^2+^ using enhanced sampling techniques. We used well-tempered meta-dynamics simulations^64,65^ to overcome the energy barriers in the transitions between OS and IS ion binding and vice versa. We computed the free energy surface for the L4 tetraloop folding and ion binding and answered the following questions: 1) What intermediate states are populated during the tetraloop folding and ion binding? 2) Does the cation’s IS or OS binding modes lead to the population of any non-native states in the folding? 3) Can the tetraloop fold and bind to Ca^2+^, and how is the stability of the folded state affected?

## Methods

### Unbiased MD Simulations

To prepare the system for well-tempered metadynamics simulations (WT-MetaD),^64,65^ we initially performed unbiased all-atom molecular dynamics simulations of the tetraloop motif using the GROMACS program (version 2018.6)^66,67^ to equilibrate the system. The tetraloop motif we used is a segment from the M-box riboswitch (PDB ID: 2QBZ)^16^ (Fig. 1B,C, S1A). The initial coordinates for the eight-nucleotide-long hairpin motif (U67-A74) are extracted from the X-ray crystal structure of the M-box riboswitch (PDB ID: 2QBZ),^16^ and we removed the phosphate group at the 5 -end. The crystal structure of the hairpin motif contains a bound Mg^2+^ ion that is IS coordinated to the O2P atom of the A72 nucleotide (designated as Mg^2+^-tetraloop system) (Fig. 1C, S1A). To assess the specific affinity of Mg^2+^ towards RNA tetraloop in comparison to the other divalent metal ions, we prepared another system by replacing the Mg^2+^ with the biologically relevant Ca^2+^ ion (designated as Ca^2+^-tetraloop system). Both the systems are solvated with 8156 TIP4P-EW^68^ water molecules in a cubic box of 6.3 nm with periodic boundary conditions in all directions. The monovalent and divalent ion concentrations are 46 mM and 7 mM, respectively, for both systems (Table 1). We used modified AMBER ff14 force field^69^ (DESRES ff) for RNA. The monovalent and divalent ion parameters are taken from ref.^70,71^

**Table 1:** Description of systems.

| System | Metal ion concentration | Tetraloop atoms | Number of $\text{K}^+$ ions | Number of divalent ions | Number of $\text{Cl}^-$ ions |
| --- | --- | --- | --- | --- | --- |
| $\text{Mg}^{2+}$-tetraloop | $[\text{K}^+] \approx 46 \text{ mM}$,<br>$[\text{Mg}^{2+}] \approx 7 \text{ mM}$ | 257 | 7 | 1 | 2 |
| $\text{Ca}^{2+}$-Tetraloop | $[\text{K}^+] \approx 46 \text{ mM}$,<br>$[\text{Ca}^{2+}] \approx 7 \text{ mM}$ | 257 | 7 | 1 | 2 |

We minimized the initial solvated structure in the presence of ions using the steepest descent minimization algorithm with a maximum force cutoff of 1000 kJ/(mol. nm). The systems are equilibrated in three consecutive steps: (i) NVT simulation of solvent molecules at temperature *T* = 300 K for 150 ns by restraining the positions of RNA atoms and ions using a harmonic potential with a force constant of 1000 kJ/(mol nm^2^), (ii) NVT simulation of solvent molecules and ions at *T* = 300 K for 150 ns by the restraining positions of RNA atoms and (iii) NPT simulation of solvent molecules and ions at *T* = 300 K and pressure *P* = 1 atm for 150 ns by restraining the positions of RNA atoms. After equilibration, we performed NPT simulation at *T* = 300 K and *P* = 1 atm with restraints on all the atoms removed. The *T* and *P* are maintained using velocity rescaling thermostat^72^ with time constant *τ*_*t*_= 0.1 ps and Parrinello-Rahman barostat^73^ with time constant *τ*_*p*_ = 2 ps. Lennard-Jones interactions are truncated at 10 Å. The electrostatic interactions are calculated using the particle-mesh Ewald (PME)^74^ method with a grid size of 1.6 Å and real space cutoff of 10 Å. All the covalent bonds containing hydrogen atoms are kept rigid using the LINCS algorithm.^75^ The equation of motion is integrated using the leapfrog integrator with a time step of 2 fs.

### Well-Tempered Meta-Dynamics Simulations

To probe the effect of metal ion IS and OS binding modes on the tetraloop folding, we performed well-tempered metadynamics simulations (WT-MetaD)^64,65^ using PLUMED 2.5.1^76,77^ and GROMACS 2018.6. We used three collective variables (CVs): (i) distance between the metal ion and phosphate oxygen O2P of nucleotide A72 in the tetraloop, *d* (Fig. S2A), (ii) coordination number, *CN* around metal ion (Fig. S2B) and (iii) *eRMSD* of the tetraloop (Fig. S2C). As the metal ion (Mg^2+^) interacts with the O2P atom of nucleotide A72 in the crystal structure, we chose *d* as one of the CVs as it can distinguish between the inner shell (IS) and outer shell (OS) binding modes of Mg^2+^ to the nucleotide A72. *CN* is calculated using the equations

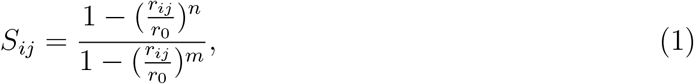

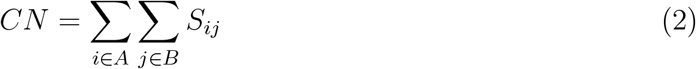

where group *A* is the metal ion (Mg^2+^ or Ca^2+^) and group *B* contains only the heavy atoms in the tetraloop binding pocket (oxygen and nitrogen atoms of C68, C69, A70, A71, A72, and G73). We did not include the atoms of water molecules in group *B*. The cutoff distance *r*_0_ = 0.50 nm and *r*_*ij*_ is the distance between atoms *i* and *j* from groups *A* and *B*, respectively. The values of *n* and *m* determine the steepness of the switching function. We have chosen the default values *n* = 6 and *m* = 12 to calculate *CN*, and it quantifies the number of contacts between atoms in groups A and B. *eRMSD* quantifies the deviation in the orientation of all the nucleobases with respect to the crystal structure.

We initiated the WT-MetaD simulations using the previously equilibrated systems from the unbiased MD simulations with the abovementioned parameters. We ran NPT simulations at *T* = 300 K and *P* = 1 atm for Mg^2+^-tetraloop (*≈* 6.5 *μ*s) and Ca^2+^-tetraloop (*≈* 4.3 *μ*s) system. Gaussian width for *eRMSD, CN*, and *d* in the WT-MetaD simulations are 0.1, 0.1, and 0.01 nm, respectively. Initial Gaussian height, deposition time, and bias factor are 0.5 kJ/mol, 1 ps, and 15, respectively. For the CV, *d*, we implemented a lower wall cut-off of 0.15 nm and an upper wall cut-off of 0.8 nm. We used sum hills and REWEIGHT BIAS modules embedded in PLUMED for reweighting simulation data. The module sum hills is used for projecting the free energy surfaces (FESs) onto the biasing CVs (*d, CN* and *eRMSD*). The module REWEIGHT BIAS is used to reweight the probability distributions and free energy surfaces onto other CVs. The convergence of WTMetaD simulations is examined by computing the free energy profile as a function of three different CVs at different simulation times. The changes in the free energy surface were minimal with increasing time, confirming the simulation’s convergence (Fig. S3). To further confirm the convergence of the simulations, we extracted the last 1.5 *μ*s (Mg^2+^-tetraloop) and 1 *μ*s (Ca^2+^-tetraloop) to estimate error in free energies of the above mentioned three CVs (*d, CN* and *eRMSD*) through block-analysis (Figure S4).

### RNA Force Field and Ion Parameters

We used the modified AMBER ff14 RNA force field^69^ (DESRES ff) as it was shown to reproduce the experimental structural and thermodynamic properties of diverse RNA structures, including single-stranded RNAs, RNA duplexes, RNA hairpins, and riboswitches. The AMBER ff14 RNA force field^69^ when used along with the TIP4P-D water model^78^ and CHARMM 22 ion parameters^79^ effectively captured native A-form conformations while avoiding artificial intercalated structures often observed in other force fields such as ff99bsc0*χ*_*OL*3_ ^80–84^ and ff99bsc0*χ*_*OL*3_+gHBfix.^62,63^ The tetraloop sequence we studied, CAAA, consists predominantly of cytosine (C) and adenine (A) nucleotides. Similar tetraloop sequences, namely AAAA, CAAU, and CCCC, were simulated using DESRES ff and demonstrated a significant population of A-form conformations (*≈* 50-80%), consistent with the experimental observations. ^69^ However, DESRES ff was shown to have limitations as it tends to overstabilize RNA helical regions, duplexes, and tetraloops.^69^ Limitations of this force field were also observed in modeling large, complex RNAs with kink-turns or L1 stacks.^62,63^

For monovalent and divalent ions, we used parameters from Joung et al.^70^ and the Li et al.,^71^ respectively, which are compatible with the TIP4P-EW water model. ^68^ Previous simulation studies^63^ have shown that for most of the tetraloop systems, the results obtained using the AMBER ff14 RNA force field^69^ are independent of the monovalent ion parameters. For the divalent ion parameters, we used the final optimized Mg^2+^ and Ca^2+^ parameters from Table 8 in Li et al.^71^ paper. Since the individual hydration-free energies (HFE) and ion-oxygen distance (IOD) parameter sets could not simultaneously reproduce both experimental HFEs and IODs accurately, we chose the optimized set, which is a compromise between the HFE and IOD sets (Table 8 in Li et al. ^71^). Parameters of the optimized set achieved *±* 2 kcal/mol accuracy of experimental relative HFE while preserving the coordination number (CN) of most divalent metal ions. Our choice of the TIP4P-EW water model will have an effect on the conformational sampling in the simulations compared to the TIP4P-D water model. However, as inferred from the previous studies, the conclusions might not be affected significantly.^62^

There are suggestions in the literature improving the parameters for the divalent ions.^71,85–88^ Studies have shown that 12-6-4 Lennard-Jones (LJ) type nonbonded potential for divalent ions, which can also take into account the charge and induced-dipole interactions, are better in balancing the ion-water and ion-RNA interactions.

### Data Analysis

*Coordination Number of Metal Ions Involved in Inner-Shell Contacts* 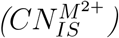. To quantify the tetraloop-ion inner-shell contact formation, we computed a modified coordination number, 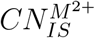, which quantifies the coordination of metal ions, M^2+^ (Mg^2+^ and Ca^2+^) with RNA heavy atoms (oxygen and nitrogen) in their first solvation shell.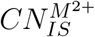 is calculated using eq. 1 and 2. In computing 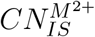, the divalent metal ion *M* is in group *A* and the RNA heavy atoms mentioned in Table S1 are present in group *B* (Fig. S5A,B). The group *B* atoms mentioned in Table S1 are observed in the first solvation shell of the metal ions in the simulation trajectories for Mg^2+^-tetraloop system and Ca^2+^-tetraloop system. We did not include the atoms of the water molecules in group *B*. We also defined a coordination number, 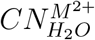 to estimate the number of water molecules present in the first solvation shell of the metal ions, M^2+^ (M = Mg, Ca). Here, the metal ion belongs to group *A*, and the oxygen atoms of all the water molecules in the simulation box belong to group *B*.

For the Mg^2+^-tetraloop system, we used *r*_0_ = 0.27 nm, *n* = 6 and *m* = 12. For the Ca^2+^-tetraloop system, we used *r*_0_ = 0.33 nm, *n* = 12 and *m* = 24. The values of *r*_0_ for Mg^2+^ and Ca^2+^ are the distance between the metal ion center and the peak of the first water solvation shell in the radial distribution function of the metal ion and water (Fig. S6). We used the default *n* and *m* values for Mg^2+^-tetraloop system. For Ca^2+^-tetraloop system we had to use larger *n* and *m* values leading to a sharp steepness in the switching function. The first solvation peak of the radial distributions function of Ca^2+^ shows a broader distribution compared to Mg^2+^ indicating that the Ca^2+^ solvation shell is less rigid due to lower charge density (Fig. S6). Hence, it was necessary to use higher values of *n* and *m* to compute 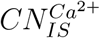 and 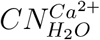. The parameters used in the calculation of *CN* and *CN* _*IS*_ are in Table S1.

#### Root Mean Square Structural Deviation (RMSD)

To quantify the structural deviation from the native structure, we computed root-mean-square deviation (*RMSD*) of the tetraloop conformations with reference to the crystal structure (Fig. 1B).

#### Radius of Gyration (R_g_)

We computed the radius of gyration (*R*_*g*_) of the tetraloop conformations using 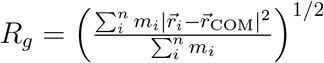, where the position of the tetraloop center of mass is given by 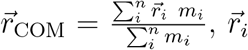 and *m*_*i*_ are the position and mass of atom *i*, and *n* is total number of atoms in the tetraloop.

## Results

### Metal Ions Form Multiple Inner Shell (IS) Contacts with the RNA Heavy Atoms

L4 tetraloop is an eight-nucleotide long hairpin motif that contains a bound Mg^2+^ ion in its crystal structure (Fig. 1B,1C,S1A,S1B). Specific binding of ions to RNA occurs through IS and OS coordination^23–27^ (Fig. 1A). In the L4 tetraloop, Mg^2+^ ion is IS coordinated to the phosphate oxygen (O2P) atom of nucleotide A72 (Fig. 1C and S1A). The other contacts of Mg^2+^ with the tetraloop are through water-mediated OS interaction by four water molecules present in the first solvation shell of Mg^2+^ (Fig. 1C and S1B). To investigate the role of these ion binding modes in specific ion sensing by RNA, we need to sample IS to OS transitions and vice versa within the simulation timescale. The Mg^2+^ IS-OS transitions are associated with high free energy barriers^29,89^ 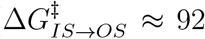 kJ/mol and 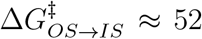 kJ/mol). We note that the exact barrier heights are sensitive to the ion force field parameters.^86,90^ However, the barrier height is *≫ k*_B_*T*, where *k*_B_ is the Boltzmann constant. Therefore, we performed well-tempered metadynamics (WT-MetaD) simulations to overcome the free energy barriers and capture all the possible ion binding modes.

We performed WT-MetaD simulations using three collective variables (CVs) to probe the tetraloop folding and ion binding (see Methods and SI): (i) distance between the metal ion (M^2+^) and phosphate oxygen O2P of nucleotide A72 (*d*) (Fig. S2A), (ii) coordination number of the metal ion (*CN*) due to its interaction with the tetraloop heavy atoms (O and N atoms) (Fig. S2B) (see SI for details), and (iii) eRMSD of the tetraloop (*eRMSD*)^55,61,91,92^ (Fig. S2C). The CV *d* monitors the formation of IS contact between the metal ion and A72-O2P atom present in the tetraloop, *CN* quantifies the coordination number around the metal ion, and *eRMSD* monitors the deviation in nucleobase’s orientation with respect to the crystal structure.

In the WT-MetaD simulations, we monitored the variation in the CVs (*d, CN*, and *eRMSD*) as a function of time. We observed: (i) multiple metal ion IS (*d <* 0.3 nm) to OS (*d >* 0.3 nm) transitions, (ii) tetraloop unfolded (*eRMSD >* 2.0) to native-like folded (*eRMSD <* 1.0) transitions, and (iii) tetraloop-metal ion bound (*CN >* 1) to unbound (*CN ≈* 0) transitions for both Mg^2+^-tetraloop and Ca^2+^-tetraloop systems (Fig. S7A,B). To identify the different states populated during these transitions in the Mg^2+^-tetraloop system, we computed the two-dimensional free energy surface (FES) *G*(*X, Y*), where *X* and *Y* ∈*{d, CN, eRMSD}* (Fig. S8). The FES *G*(*eRMSD, CN*) shows that multiple intermediate states are populated (Fig. S8A). Although the native-like folded state (*eRMSD ≈* 0.8) with *CN ≈* 10 has a significant population, the most populated states are present within *eRMSD >* 1.5 and 3 *< CN <* 12. Interestingly, intermediate (*eRMSD ≈* 1.5) and unfolded (*eRMSD ≈* 2.5) conformations form higher coordination (5 ≤ *CN* ≤ 10) with the metal ions compared to the folded states (*eRMSD ≈* 1). Further FES *G*(*eRMSD, d*) shows that IS-coordinated states (*d <* 0.3 nm) are formed mainly by intermediate and unfolded tetraloop conformations compared to the folded conformations (Fig. S8B). The higher propensity of partially folded intermediates and unfolded conformations of the tetraloop to form high coordination structures with the metal ions suggests that the structural features of the tetraloop play a critical role in metal ion sensing.

*G*(*CN, d*) also shows that IS-coordinated Mg^2+^ with A72 nucleotide (*d ≈* 0.2 nm) have coordination between 4 to 11 (Fig. S8C). The OS-coordinated Mg^2+^ (*d ≈* 0.4 nm) mostly populates structures with *CN ≈* 3 (basin 1) and 10 (basin 2) (Fig. S8C and S9A). However, OS-coordinated structures with high *CN ≈* 10 are energetically unfavorable. To understand the origin of high *CN*, we extracted the structures from basins 1 and 2. We discovered that even though Mg^2+^ is OS-coordinated with A72-O2P atom (*d ≈* 0.4 nm), it can form multiple conformations where Mg^2+^ is IS-coordinated (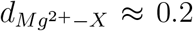 nm) with other electronegative atoms, X = A72-O1P, C69-O2, G71-O2P etc (non-native ion-tetraloop IS contacts) (Fig. S9C,D). To further confirm Mg^2+^ IS-coordination with nucleotides other than A72, we computed the coordination number around Mg^2+^ ion considering only the tetraloop atoms present in the first solvation shell of the metal ion, 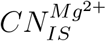 (see Data Analysis section) for basin 1 and 2 structures (Fig. S9B). We observed that basin 1 (basin 2) structures form one (three) IS-coordination with nucleotides other than A72-O2P. There are multiple events throughout the simulation where Mg^2+^ is OS-coordinated with A72-O2P and IS-coordinated with any of the eight electronegative atoms mentioned in Table S1 (Fig. S10A and S5A). The FES projected onto the distance between the metal ion and any of the eight IS-coordinated atoms also confirms that there is a significant probability of forming IS coordination (*d <* 0.3 nm) between metal ion and tetraloop atoms (Fig. S10B). The decrease in the number of wa-ter molecules 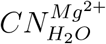 (see Data Analysis section) and increase in the number of tetraloop atoms 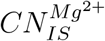 around Mg^2+^ in the first solvation shell confirms that the IS-coordinated structures are formed through the removal of water from the first solvation shell of the metal ion (Fig. S11A). These observations indicate that even though the Mg^2+^ ion is bound to the A72-O2P atom through IS coordination in the folded state, it can form multiple IS contacts with the other electronegative atoms in the tetraloop during its folding. Additionally, the structural features of the tetraloop play a critical role in forming these IS-coordinated structures.

### Tetraloop Folding Free Energy Surface Shows Population of Multiple Inter-mediate Structures

In the previous section, we showed that metal ions can form IS contacts with different RNA atoms, and the conformations of the tetraloop plays a vital role in facilitating these IS contacts. To understand the role of tetraloop structure in metal ion sensing, we probed IS contact formation by computing 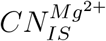 and *RMSD* (see Data Analysis section in the Methods in SI). Although *eRMSD* is generally used to characterize RNA structures, it only considers the orientation of nucleobases and cannot capture the structural deviation of the RNA backbone. Therefore, we used *RMSD* to characterize the structural deviation of the tetraloop backbone and nucleobase orientation relative to the crystal structure.

We computed the two-dimensional FES by reweighing the simulation trajectories onto the 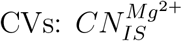 and *RMSD* using REWEIGHT BIAS module embedded in PLUMED^93^ (Fig. 2). The FES shows that the tetraloop populates multiple partially folded intermediate states. There are five minima corresponding to 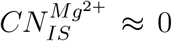 (basin **IS**_**0**_), 1 (basin **IS**_**1**_), 2 (basin **IS**_**2**_), 3 (basin **IS**_**3**_) and 4 (basin **IS**_**4**_). Basins **IS**_**0**_ to **IS**_**4**_ populate the tetraloop structures where the metal ion forms 0 to 4 IS contacts (Fig. S7C). The remaining coordination of Mg^2+^ is fulfilled by the water molecules. Although Mg^2+^ can technically form six IS contacts, we observed a maximum of four IS contacts (Fig 2, snapshot **IS**_**4**_(I1)). The structural constraints in the tetraloop prevents the formation of six IS contacts.

**Figure 2.**
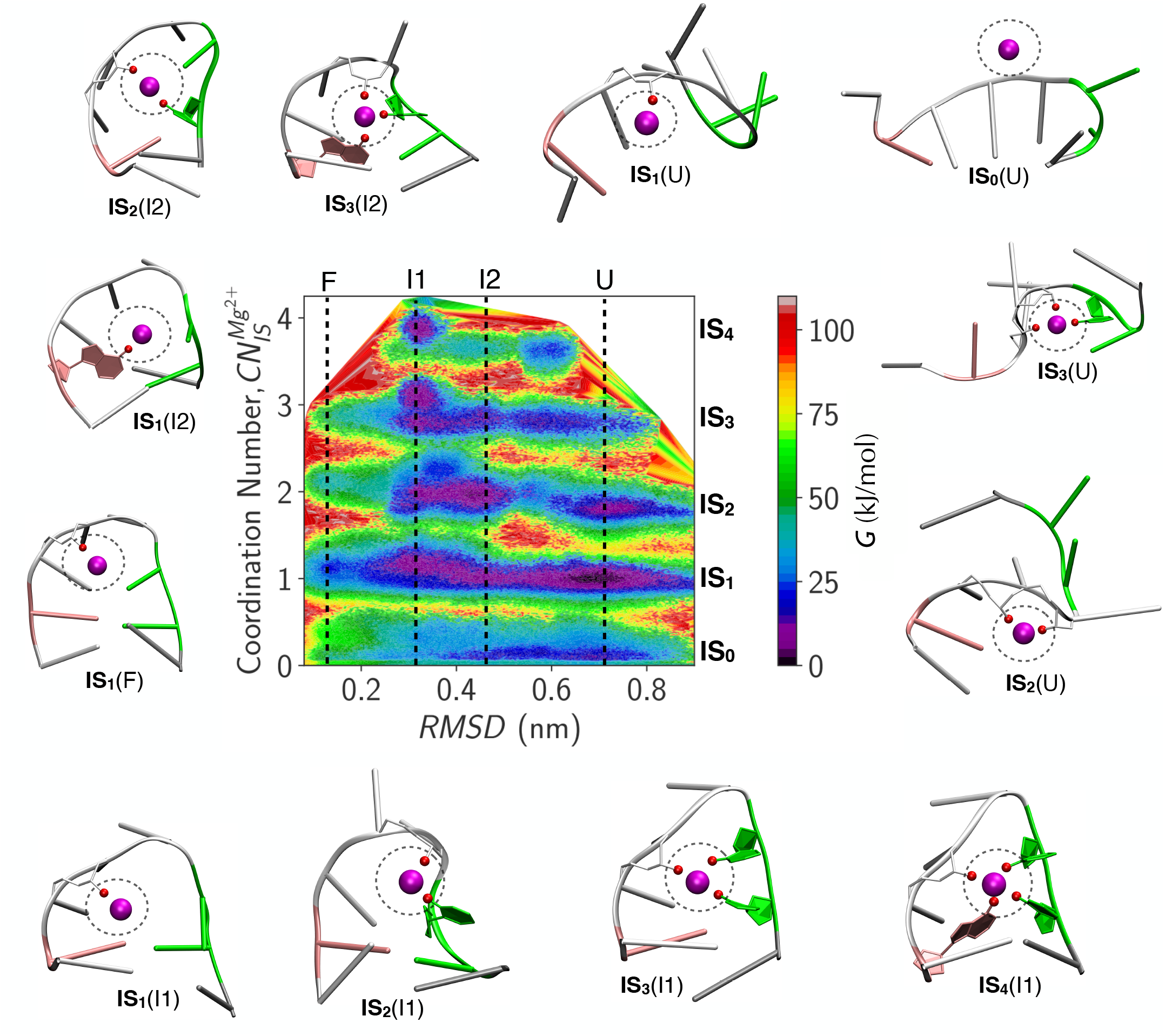
FES of the L4 tetraloop projected onto the CVs, 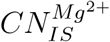 and *RMSD*. The color bar represents free energy, Δ*G*. Five distinct basins (**IS**_**0**_ to **IS**_**4**_) are populated based on 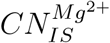 (0 to 4). Based on *RMSD*, tetraloop structures are categorized as folded (F), intermediate1 (I1), intermediate2 (I2) and unfolded (U), which are shown with the black dotted lines. Representative structures corresponding to the basins are shown. The Mg^2+^ ion and RNA atoms involved in IS coordination are shown with magenta and red spheres, respectively. The black circle highlights the number of inner-shell interactions.

The tetraloop adopts a range of conformations between folded (*RMSD ≈* 0 nm) and unfolded (*RMSD ≈* 1 nm) states (Fig. 2). The FES projected onto *RMSD* showed four distinct minima that correspond to folded (F) (*RMSD ≈* 0.14 nm), intermediate1 (I1) (*RMSD ≈* 0.32 nm), intermediate2 (I2) (*RMSD ≈* 0.44 nm) and unfolded (U) (*RMSD ≈* 0.71 nm) states (Fig. S12A). The folded state of the tetraloop is less stable compared to the intermediates and unfolded structures as the tetraloop is a truncated part of the M-box riboswitch (Fig. S12A). In the M-box riboswitch (PDB ID: 2QBZ),^16^ the L4 tetraloop forms tertiary interactions with other parts of the structure (Fig. 1B). A70 interacts with adjacent helices through both stacking and hydrogen bonding interactions. Additionally, nucleotides C68, A71, A72, G73, and A74 form hydrogen bonds with the adjacent helices, and C67 interacts with a bridging K^+^ ion through IS coordination. These tertiary interactions are crucial in stabilizing the folded state and are absent in the truncated L4 tetraloop.

The states (F, I1, I2, and U) are not equally populated in basins with the same coordination number (basins labeled **IS**_**0**_ to **IS**_**4**_) (Fig. 2). For basins labeled 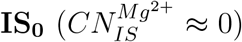 the I1, I2, and U states are more populated compared to the F state. The states F and U are absent in basins labeled 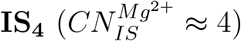, indicating that these two states are unable to form four IS contacts with Mg^2+^. In the other basins (**IS**_**1**_, **IS**_**2**_, **IS**_**3**_), states from F to U have a significant population. Therefore, the population difference in various states (folded or unfolded) of the tetraloop in basins from **IS**_**0**_ to **IS**_**4**_ further confirms that the formation of IS contacts depends on the tetraloop conformation. As a result, the free energy barriers for transitions between these five basins (**IS**_**0**_ → **IS**_**1**_, **IS**_**1**_ → **IS**_**2**_, **IS**_**2**_ → **IS**_**3**_, and **IS**_**3**_ → **IS**_**4**_) will also depend on the tetraloop conformations, i.e., whether it is folded, unfolded, or in the intermediate states (Fig. 2). Finally, we conclude that the FES for the tetraloop folding and metal ion sensing is multidimensional as Mg^2+^ can form multiple IS contacts with RNA atoms simultaneously, resulting in the population of multiple partially folded intermediate states.

The temperature jump kinetics experiments on non-GNRA sequence (cgGCUUcg) revealed two distinct intermediate states in addition to the folded and unfolded states. ^94^ The first intermediate state referred to as ensemble E features a native loop structure with an incomplete stem structure. The second intermediate state referred to as ensemble S has a non-native loop structure with a complete stem structure. We observed that the L4 tetraloop also samples conformations similar to the E-type structures in the basins **IS**_**0**_ and **IS**_**1**_ for both I1 and I2 states (Fig. S13). The L4 tetraloop conformations similar to the S-type structures are observed in the basins **IS**_**0**_, **IS**_**1**_, and **IS**_**2**_ for the I1 state.

### Free Energy Barrier for Transitions Between Different IS Contacts Depends on the Compactness of the Tetraloop Conformations

The number of IS contacts between the metal ion and tetraloop strongly depends on the tetraloop conformation. To probe the role of tetraloop conformations in metal ion sensing, we extracted conformations from the minima of various basins in the FES projected on *RMSD* (Fig. S12A) that correspond to: (i) folded (F) (*RMSD ≈* 0.14 *±* 0.01 nm), (ii) intermediate1 (I1) (*RMSD ≈* 0.32 ± 0.01 nm), (iii) intermediate2 (I2) (*RMSD ≈* 0.44 *±* 0.01 nm), and (iv) unfolded (U) (*RMSD ≈* 0.71 *±* 0.01 nm) states (shown with black dashed lines in Fig. 2). For each state (F, I1, I2, U), we projected the FES on 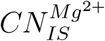 to quantify the stability of different basins (**IS**_**0**_ to **IS**_**4**_) and the transition barrier between these basins (Fig. 3A and S14).

**Figure 3.**
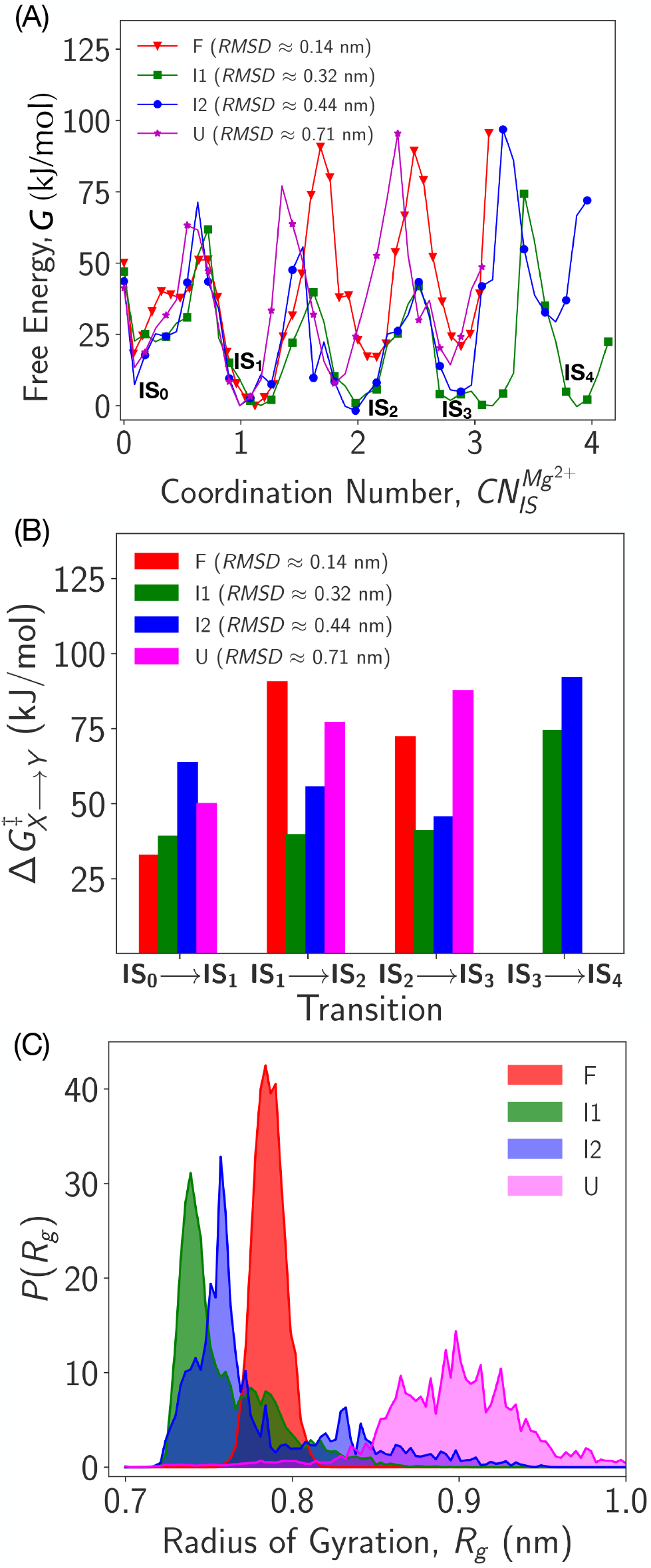
(A) FES projected onto 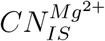 for different L4 tetraloop states (F, I1, I2 and U). Basins **IS**_**0**_ to **IS**_**4**_ corresponds to the structures with 0 to 4 IS contacts between Mg^2+^ and tetraloop. (B) The free energy barrier, 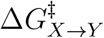 associated with the transition from basin *X* (= **IS**_**0**_, **IS**_**1**_, **IS**_**2**_, **IS**_**3**_) to *Y* (= **IS**_**1**_, **IS**_**2**_, **IS**_**3**_, **IS**_**4**_) for F (red), I1 (green), I2 (blue) and U (magenta) states. (C) Probability distribution of *R*_*g*_ for states F, I1, I2, and U.

The FES projected onto 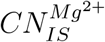 shows five distinct minima associated with 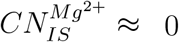 (basin **IS**_**0**_) to 4 (basin **IS**_**4**_) (Fig. 3A). Basin 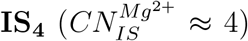 is absent for F and U states as the structural constraints in the folded and unfolded conformations prevent it to form four IS contacts with the Mg^2+^ ion. The relative stability of various basins (**IS**_**0**_ to **IS**_**4**_) differs for various states (F to U). For folded state conformations, the relative stability follows the trend: 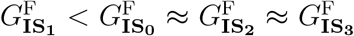, which indicates that the formation of single IS contact is more facile in the F state compared to zero or more than one IS contacts. A similar trend is also observed for unfolded states where the population of basin **IS**_**1**_ with one IS contact is more stable than other basins, i.e., 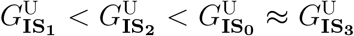. Interestingly, I1 and I2 state conformations prefer to form more than one IS contact. For I1 state, relative stability follows: 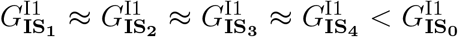. For the I2 state, the basins **IS**_**1**_, **IS**_**2**_ and **IS**_**3**_ are more stable than basins **IS**_**0**_ and **IS**_**4**_, i.e. 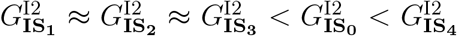.Therefore, the relative stability of various basins (**IS**_**0**_ to **IS**_**4**_) strongly depends on the tetraloop conformation. As a result, the RNA conformations could play a crucial role in overcoming the free energy barrier for the transition from one basin to another with different IS contacts.

To gain insights into the role of RNA conformations (F, I1, I2 and U) on the transitions between basins with different IS coordinated structures (**IS**_**0**_ to **IS**_**4**_), we computed the free energy barrier associated with these transitions, i.e. 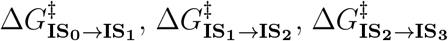 and 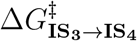 (Fig. 3B). The free energy barrier for the transition from basin 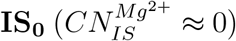 to **IS**_**1**_ 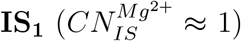 follow the trend: Δ*G*^‡,F^ *<* Δ*G*^‡,I1^ *<* Δ*G*^‡,U^ *<* Δ*G*^‡,I2^, which indicates that F and I1 conformations prefer to form single IS coordination with Mg^2+^ compared to the U and I2 structures. Also, among the F conformations, the **IS**_**1**_ state has the lower free energy (Fig. 3A), indicating that among the conformations with single IS contact, the folded state is the most stable and forms faster due to lower free energy barriers supporting the formation of crystal-like structure (Fig. 1C, S1A).

For the transitions between basin **IS**_**1**_ to **IS**_**2**_ and **IS**_**2**_ to **IS**_**3**_, the intermediate structures I1 and I2 have lower barriers compared to the F and U conformations. For the **IS**_**1**_ to **IS**_**2**_ transition, the free energy barrier follows Δ*G*^‡,I1^ *<* Δ*G*^‡,I2^ *<* Δ*G*^‡,U^ *<* Δ*G*^‡,F^. For the **IS**_**2**_ to **IS**_**3**_ transition, the order is Δ*G*^‡,I1^ *<* Δ*G*^‡,I2^ *<* Δ*G*^‡,F^ *<* Δ*G*^‡,U^. The free energy barrier for transitioning from basin **IS**_**3**_ to **IS**_**4**_ is high in both I1 and I2 states, and follows the order: Δ*G*^‡,I1^ *<* Δ*G*^‡,I2^.

We hypothesized that the origin of the different free energy barriers for transitions between states depends on the compactness of the tetraloop conformations. To test the hypothesis, we computed the probability distribution of the tetraloop radius of gyration (*R*_*g*_) (see Methods), *P* (*R*_*g*_), which shows that the intermediate conformations (I1 and I2) are more compact compared to the F and U states (Fig. 3C). The folded (F) state has a narrow distribution of *R*_*g*_ centered around *R*_*g*_ *≈* 0.77 nm. Unfolded conformations have a broad size distribution ranging from *R*_*g*_ = 0.7 to 1 nm. The I2 state is more compact compared to the F and U states and has a maximum population around *R*_*g*_ *≈* 0.75 nm. However, the I1 state is most compact with a significant population at *R*_*g*_ *≈* 0.72 nm. The size quantified using *R*_*g*_ follows the order: I1 *<* I2 *<* F *<* U. The *R*_*g*_ distribution shows that the smaller the size of the tetraloop, the higher the propensity to form multiple IS contact formation. As a result, I1 and I2 conformations are more prone to form multiple IS contacts 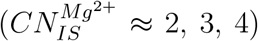 compared to F and U states. The folded RNA state easily forms a single IS contact (basin **IS**_**1**_) because it is structurally rigid, preventing reorganization of their backbone/nucleobases for multiple IS contacts (basins **IS**_**2**_, **IS**_**3**_, and **IS**_**4**_). Higher compaction in the I1 state (lower *R*_*g*_ conformations) facilitates the population of tetraloop conformations with three (basin **IS**_**3**_) and four IS contacts (basin **IS**_**4**_) compared to the I2 state with relatively larger R_*g*_ conformations. Unfolded conformations with higher *R*_*g*_ cannot facilitate establishing multiple IS contacts with the ion. Therefore, the flexibility of I1 and I2 states with higher RNA compaction facilitates the formation of multiple simultaneous IS contacts compared to F and U states.

### Population of Non-native Tetraloop-Ion Inner-Shell Bound Structures Contribute to the High Energy Barrier During L4 Tetraloop Folding to its Native Structure

Tetraloop folding and metal ion binding is a concerted process during which the phosphate oxygen atom of A72 in the tetraloop coordinates with the metal ion through OS coordination and transitions to an IS coordination via the elimination of a water molecule from its first solvation shell to the second solvation shell (OS-to-IS transition) and vice versa.^29^ This process is known as the “RNA-water exchange process” (Fig. 4A). To understand the free energy barriers involved in the RNA-water exchange process, we computed the FES projected onto 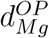 (distance between Mg^2+^ and O2P atom of A72) and compared it to the FES of the water-water exchange process^95^ (Fig. 4A) projected onto 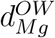 (distance between Mg^2+^ and oxygen atom of water molecule) (Fig. 4B).

**Figure 4.**
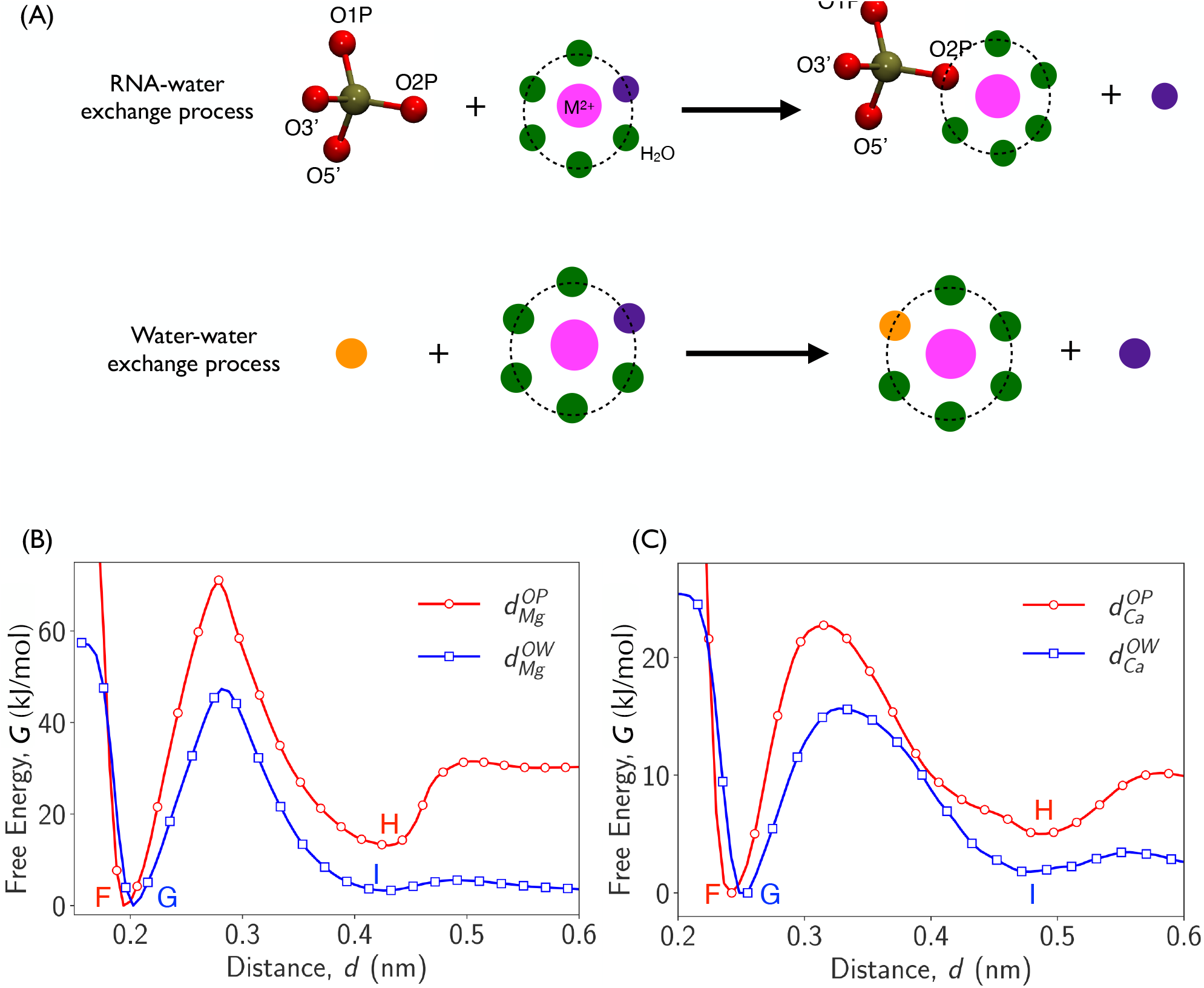
(A) Schematic diagram describing RNA-water exchange process (RNA-ion binding). Phosphate oxygen (O1P) atoms of RNA are represented with red spheres. In this process, OP1 atom of RNA participates in direct ion binding in the first solvation shell (shown with dotted circle) by eliminating one water molecule (violet bead) from the first solvation shell. Metal ion and water molecules are shown with magenta and green beads, respectively. Schematic diagram describing water-water exchange process (water-ion binding) where one water molecule (yellow bead) present in the second solvation shell of the metal ion replaces another water molecule (violet bead) present in the first solvation shell of metal ion.(B) FES projected on the distance between Mg^2+^ and water, 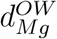 (blue line with squares) or Mg^2+^ and RNA, 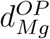 (red line with circles). Basin F (G) and H (I) are the IS and OS-bound states in the RNA-ion (water-ion) binding process. (C) Same for Ca^2+^.

In the case of the water-water exchange process with Mg^2+^ ion (Fig. 4A), as expected, the IS (basin G) and OS-coordinated (basin I) are equally stabilized as there is no change in the water solvation of the Mg^2+^ ion at the end of the exchange process (Fig. 4B, S15H). The free energy barrier for OS-to-IS or IS-to-OS transition process for Mg^2+^ is *≈* 47 kJ/mol (Fig. 4B). However, the stability and free energy barrier for the RNA-water exchange process is significantly different from the water-water exchange process. The stability of native tetraloop-ion IS structure (basin F) is more stable compared to its OS structure (basin H) because the IS interaction between Mg^2+^ and the phosphate oxygen atom of A72 is stronger compared to the interaction between Mg^2+^ and water (Fig. 4B). The free energy barrier for IS-OS/OS-IS transition is higher for the RNA-water exchange process compared to the water-water exchange process as relatively stronger non-native Mg^2+^-tetraloop IS contacts should be broken compared to the Mg^2+^-water contacts (Fig. S15A-G).

### Ca^2+^ Cannot Stabilize the Loop Closing Interactions in the Tetraloop Due to Lower Charge Density

To further gain insights into the metal ion sensing by RNA, we compared our observations for Mg^2+^ binding to the tetraloop with Ca^2+^, another biologically relevant divalent metal ion. We replaced Mg^2+^ with Ca^2+^ ion in the simulation box and performed WT-MetaD simulations using the same protocols used for the Mg^2+^-tetraloop system. Since Ca^2+^ ion (radius *R* = 1.657 Å) has a lower charge density compared to Mg^2+^ ion (*R* = 1.353 Å), the barrier for OS to IS transition for Ca^2+^ ion (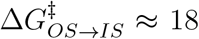 kJ/mol and 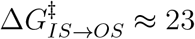 kJ/mol) is lower compared to Mg^2+^ ion (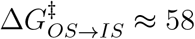 kJ/mol and 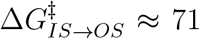 kJ/mol) (Fig. 4B,C). The radius of the ions are taken from the force field.^71^ Although the free energy barriers are sensitive to the force field parameters of the ions, experiments^96–98^ also show that the water exchange rate from the first solvation shell of Ca^2+^ is almost three orders faster compared to Mg^2+^. In the simulations, we observe more transitions between IS-to-OS or OS-to-IS for Ca^2+^ compared to Mg^2+^ due to the low transition energy barriers for Ca^2+^ (Fig. S7A,B).

Similar to the observations of the Mg^2+^-tetraloop system, the FES *G*(*eRMSD, CN*) shows that intermediate and unfolded (*eRMSD >* 2) tetraloop conformations are more prone to bind with Ca^2+^ (2 *< CN* ≤ 10) compared to folded (*eRMSD* ≤ 1) conformations (Fig. S8D). *G*(*eRMSD, d*) further confirms that intermediate and unfolded states (*eRMSD >* 2) form both IS (*d <* 0.3 nm) and OS (*d* ≥ 0.3 nm) contacts with Ca^2+^ (Fig. S8E). *G*(*CN, d*) shows that the OS-coordinated structures populate multiple minima around *CN ≈* 3, 6, 8 for Ca^2+^ similar to Mg^2+^ (Fig. S8F).

We also computed the FES for Ca^2+^ using the CVs, 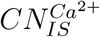 and *RMSD* to understand Ca^2+^ sensing by the tetraloop (Fig. 5A) (see Methods). We observed seven distinct minima corresponding to 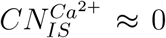 *≈* 0 (basin **IS**_**0**_), 1 (basin **IS**_**1**_), 2 (basin **IS**_**2**_), 3 (basin **IS**_**3**_), 4 (basin **IS**_**4**_), 5 (basin **IS**_**5**_), 6 (basin **IS**_**6**_) and 7 (basin **IS**_**7**_), which indicates that Ca^2+^ can form 0 to 7 IS contacts with the tetraloop due to its larger size (Fig. 5A and S7D). Based on *RMSD*, we classified tetraloop conformations belong to the following states: folded (F) (*RMSD ≈* 0.14 nm), intermediate1 (I1) (*RMSD ≈* 0.34 nm), intermediate2 (I2) (*RMSD ≈* 0.58 nm), unfolded1 (U1) (*RMSD ≈* 0.68 nm) and unfolded2 (U2) (*RMSD ≈* 0.80 nm)

**Figure 5.**
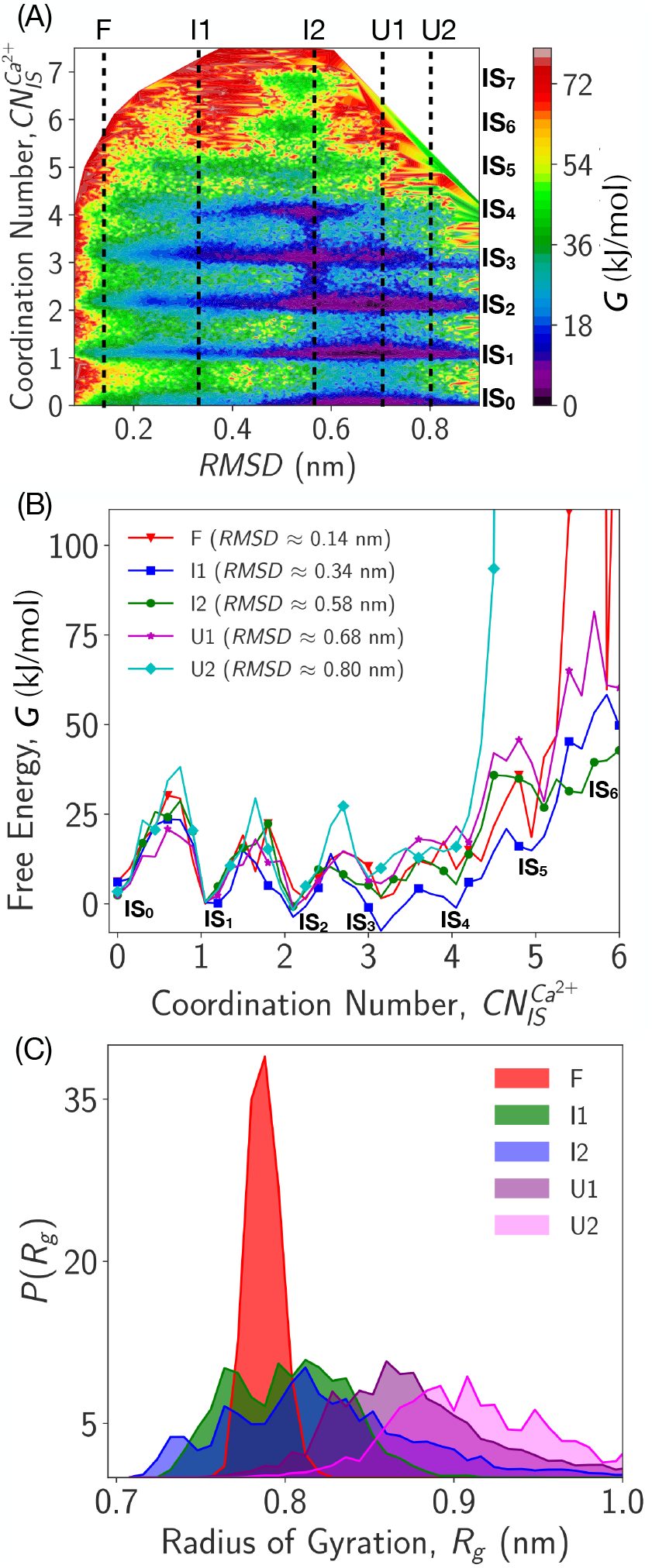
(A) FES of the L4 tetraloop projected onto the coordination number around metal ion Ca^2+^ due to inner-shell (IS) contact formation with RNA atoms, 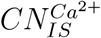 and *RMSD* from crystal structure (PDB ID: 2QBZ), *RMSD* in the presence of Ca^2+^ ion. The color bar represents free energy, *G*. Eight distinct basins (**IS**_**0**_ to **IS**_**7**_) are populated based on the 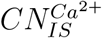 (0 to 7). Based on *RMSD*, tetraloop structures are categorized as folded (F), intermediate1 (I1), intermediate2 (I2), unfolded1 (U1) and unfolded2 (U2) which is shown with the black dotted line. (B) FES projected on 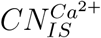 is shown for different L4 tetraloop states: F, I1, I2, U1, and U2. (C) Probability distribution of *R*_*g*_ for state F, I1, I2, U1 and U2 of L4 tetraloop in the presence of Ca^2+^.

(Fig. 5A, S12B and S16).

The FES shows that the I2, U1, and U2 states are more populated, and the F state is the least populated (Fig. 5A). The stability of the tetraloop folded state is lower in the presence of Ca^2+^ compared to Mg^2+^. Due to the low charge density of Ca^2+^, it cannot hold the loop-closing nucleotides A72 and C69 together and stabilize the tetraloop’s native-like state (Fig. 1B and S1A,B). The interaction of Ca^2+^ with the tetraloop heavy electronegative atoms is weaker compared to Mg^2+^ due to its low charge density, which is consistent with the observations from the simulations on self-splicing group-II intron. ^45^ Since Ca^2+^ ion is large, it can have a coordination number eight within its first solvation shell (*R* = 1.657 Å). Ca^2+^ can have a maximum of seven inner-shell (IS) coordinations with tetraloop atoms (Fig. S11B) and the eighth interaction is with a water molecule. In contrast, Mg^2+^, with its smaller size (*R* = 1.353 Å), can have four IS contacts with the tetraloop atoms, and water molecules fulfill the remaining two coordinations. The larger size of Ca^2+^ leading to lower charge density results in the decreased stabilization of tetraloop folded states and increased IS contact formation compared to Mg^2+^.

To understand the role of tetraloop conformations (F, I1, I2, U1, and U2 states) in populating structures with different IS contacts (**IS**_**0**_ to **IS**_**7**_), we projected the FES onto 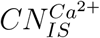 for different states (Fig. 5B). The population of basin **IS**_**7**_ is minimal, and only the I2 state, which populates the most compact conformations, can form seven IS contacts (Fig. 5A,C). Since the folded conformations (F state) are rigid, and the unfolded states (U1 and U2 states) are extended, they cannot populate structures with more than 5 IS contacts (Fig. 5A,B). Thus, similar to the Mg^2+^-tetraloop system, states with compact conformations (I1 and I2) can populate structures with IS contacts more than five, and rigid folded state (F) and the extended unfolded states (U1 and U2) populate structures with IS contacts less than five. The transition barriers for going from **IS**_**0**_ to **IS**_**7**_ in the Ca^2+^-tetraloop system are much lower than the Mg^2+^-tetraloop system (Fig. S17). Similar to the Mg^2+^-tetraloop system, the free energy barrier for IS-OS/OS-IS transition for Ca^2+^-tetraloop system is higher compared to water-water exchange process due to presence of non-native Ca^2+^-tetraloop IS contacts present in both the F and H basins. (Fig. 4C).

### Binding Specificity of Metal Ions is Higher for the Phosphate and Base Oxygen Atoms than Sugar Oxygen and Base Nitrogen Atoms

The FES of the tetraloop shows that structural compaction dictates the number of IS contacts the tetraloop can form with the ion (Fig. 2 and 5A). We further probed the propensity of IS contact formation of various electronegative atoms such as phosphate oxygen O1P and O2P, base oxygen O2 and O6, sugar oxygen O2*’*, O3*’* and base nitrogen N3, N7 with the metal ions (Fig. S5).

To quantify the propensity of the electronegative atoms in the tetraloop to form IS contacts, we computed the probability distribution of the distance between the ion and the electronegative atoms (Fig. S18, S19 and S20). From the probability distributions, we infer that both Mg^2+^ and Ca^2+^ ions have higher specificity to form IS contacts with the phosphate oxygen and base oxygen compared to sugar oxygen and base nitrogen because the backbone phosphate oxygens are negatively charged. The orientation of the bases in the tetraloop significantly influences the formation of different intermediate states, making base oxygens crucial for forming IS contacts with metal ions compared to sugar oxygens. Further, base oxygens have a higher propensity for IS contact formation than base nitrogens due to the higher electronegativity of oxygen compared to nitrogen.

The above observation where Mg^2+^ is dominantly in IS contact with the phosphate and base oxygens is in agreement with the conclusions obtained from the analysis of the RNA crystal structures. ^99–101^ These studies further establish that Mg^2+^ favors IS interaction with the base oxygens of nucleotides in the order G *>* U *>* C, and the IS interaction of Mg^2+^ with the sugar oxygens and base nitrogens are rare.^100,101^

Multiple MD studies on various RNA systems using different force fields, water models, and Mg^2+^ parameters support the conclusion that backbone phosphate oxygens have a dominant propensity for IS interaction with the Mg^2+^, followed by the base oxygens.^30,85,102,103^ However, as expected, there are quantitative differences in the propensities depending on the RNA force field and Mg^2+^ ion parameters. Simulations also show that among the phosphate oxygen atoms, O2P has a higher affinity over O1P for IS bonding with Mg^2+^.^30,102^ Significant differences were observed between the studies^30,85,102^ for the IS binding affinities of Mg^2+^ with the sugar oxygens and base nitrogens, probably due to the force field issues and these interactions were predicted to be rare from the analysis of crystal structures. ^99–101^ Development of new Mg^2+^ force fields following the guidelines prescribed by Casalino et al. ^85^ could mitigate some of inconsistencies observed in the MD simulations.

## Conclusion

We studied the folding of a tetraloop-like segment from the L4 loop of the M-box riboswitch. Interestingly, this tetraloop binds to a single Mg^2+^ ion, which makes it feasible to probe the binding mechanism of the metal ion and its role in the tetraloop folding. We show that the folding free energy surface of this small RNA segment forming the tetraloop is multidimensional, with the population of multiple intermediates categorized based on the number of inner-shell contacts the metal ion forms with the tetraloop. In most of these intermediates, the metal ion forms non-native inner shell contacts with the electronegative atoms in the tetraloop. The conformations where multiple inner shell contacts between the tetraloop and the ion are observed are more compact than the native tetraloop state. For the RNA segment to fold to the native state, all the non-native ion-tetraloop inner shell contacts with the metal ion should be disrupted by thermal fluctuations, leading to higher folding energy barriers. We further show that Ca^2+^ is less effective in stabilizing the native state of the tetraloop as its interaction with the loop closing nucleotides is weaker due to its low charge density compared to Mg^2+^.

The predictions from the simulations that Mg^2+^ forms multiple non-native inner shell contacts with various atoms in the L4 tetraloop before going into the native-like binding state can be verified by the NMR experiments. The system studied in this work, the eight nucleotide RNA segment, which folds into a tetraloop-like structure and can specifically bind to a single Mg^2+^, is probably the simplest system to probe ion-sensing by RNA experimentally. Since the system binds to only a single ion, it eliminates the complexity in the data analysis of large RNA systems, and isotope labeling of the nucleotides can help in precisely identifying where the ion interacts during the folding of this segment.

The results from this study have implications for understanding the mechanism of specific ion sensing by riboswitches in bacteria to maintain ion homeostasis. In the past decade, riboswitches that specifically bind to divalent metal ions such as Mg^2+^, Mn^2+^, Ni^2+^ and Co^2+^ and regulate their concentration have been discovered. It is essential to understand the various RNA-related structural factors and ion-related properties that contribute to the selectivity of riboswitches in binding to the ions. The results from this work show that the ions, when exploring the binding pocket in the riboswitch, can form non-native inner shell contacts with the nucleotides forming the binding pocket. These non-native inner shell contacts must be disrupted by thermal fluctuations for the ion to bind, contributing to the delay in the ion binding process. It would be interesting to explore if these non-native inner shell contacts have any role in the selective ion binding by the riboswitches.

## Supporting Information

Parameters used to calculate coordination number CVs (Table S1); Snapshots of IS and OS coordination of metal ion (Fig. S1); Schematic of CVs (Fig. S2); Convergence of WT-MetaD simulations (Fig. S3); Error bars of CVs (Fig. S4); IS contact forming tetraloop atoms (Fig. S5); RDF of ion-water (Fig. S6); CVs as a function of time (Fig S7); FES of tetraloop (Fig. S8); FES of Mg^2+^-tetraloop system (Fig. S9); Time evolution and probability distribution function (PDF) of IS contact distances (Fig. S10); Trajectories showing IS contacts formation by replacing water (Fig. S11); FES projected onto *RMSD* (Fig. S12); Experimentally observed intermediates (Fig. S13); Representative conformations for non-native IS contacts and different basins (Fig. S14 and S16); Representative conformations for non-native IS contacts and different basins (Fig. S15); Transition barriers for Ca^2+^-tetraloop system (Fig. S17); PDF of the distance between the metal ion and IS contact forming tetraloop atoms (Fig. S18, S19 and S20).

## Conflicts of Interest

There are no conflicts to declare.

## Acknowledgement

GR acknowledges funding from the National Supercomputing Mission (MeitY/R&D/HPC/2(1)/2014). SH and LB acknowledge the research fellowship from the Prime Minister’s Research Fellows (PMRF) scheme. We acknowledge the National Supercomputing Mission (NSM) for providing computing resources of “Param Pravega” at IISc and “Param Brahma” at IISER Pune, which are supported by the Ministry of Electronics and Information Technology (MeitY) and Department of Science and Technology (DST), Government of India.

## For Table of Contents Use Only

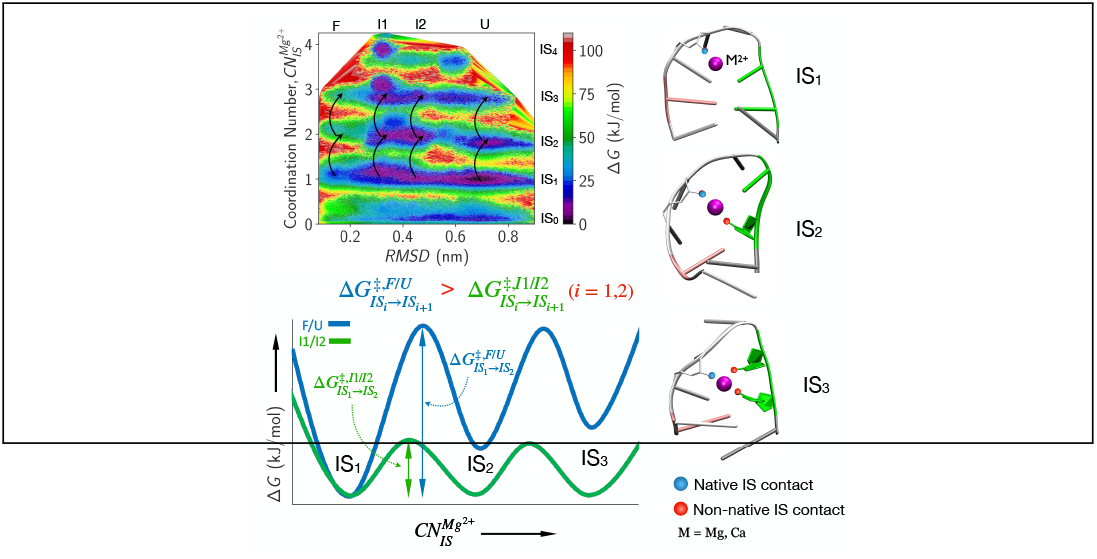

## Notes

### Competing Interest Statement

The authors have declared no competing interest.

