## Supporting Information for "Metal Ion Sensing by Tetraloop-Like RNA Fragment: Role of Compact Intermediates with Non-Native Metal Ion-RNA Inner Shell Contacts"

**Table S1: Parameters used to calculate the coordination number collective variables**

| Collective Variable | Cut-off distance ($r_0$) | Tetraloop heavy atoms used in the calculation |
|---|---|---|
| $CN$ | 0.5 nm for $Mg^{2+}$ and $Ca^{2+}$ | N and O atoms present in nucleotides C68, C69, A70, A71, A72 and G73 |
| $CN_{IS}^{Mg^{2+}}$ | 0.27 nm | A72-O2P, A72-O1P, A71-O2P, G73-O6, C69-O2, C68-N3, C68-O2, C69-N3, C69-O2′ |
| $CN_{IS}^{Ca^{2+}}$ | 0.33 nm | U67-(O5′,O4,O2,O2′,O3′), C68-(O1P,O2P,O2,N3,O2′,O3′,O5′), C69-(O1P,O2P,N3,O5′,O4′,O2,O2′,O3′), A70-(O1P,O2P,O5′,N7,O2′,O3′), A71-(O1P,O2P,N7,N1,N6,O2′,O3′,O5′,O4′), A72-(O1P,O2P,N7,N6,N1,O2′,O3′,O5′,O4′), G73-(O1P,O2P,N7,O6,N2,N3,O5′,O4′), A74-(O2P,N7,N6,N1,O2′,O3′) |

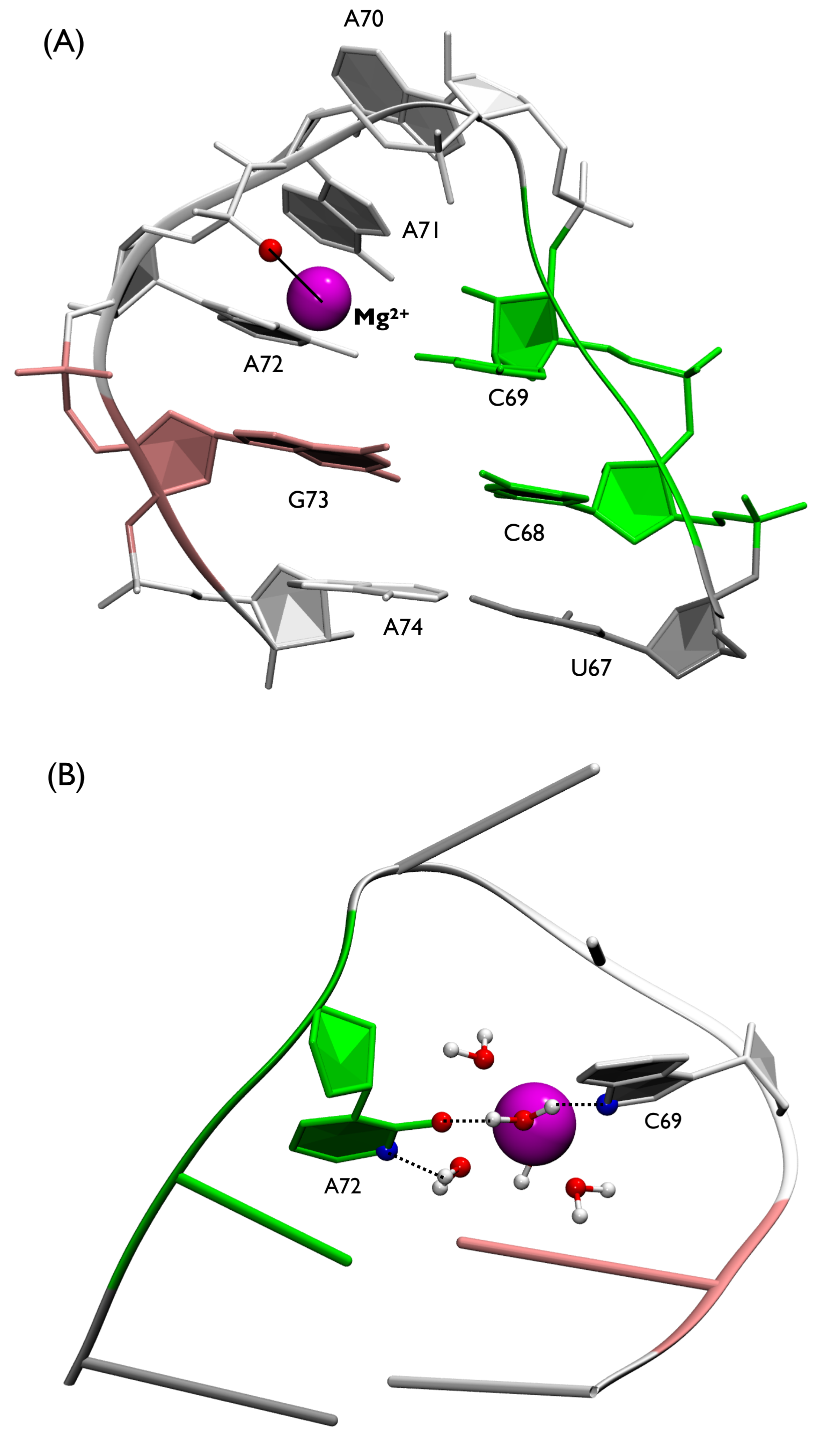


Figure S1: (A) Eight nucleotides long L4 tetraloop where the bound $Mg^{2+}$ interacts with the phosphate oxygen (O2P) atom of nucleotide A72 through IS contact. $Mg^{2+}$ is shown as a magenta sphere. The IS contact between the metal ion and the O2P atom in A72 is shown with a black solid line. (B) $M^{2+}$ ion (magenta colour, M=Mg,Ca) forms loop-closing water-mediated OS coordinations (dotted line) with base oxygens (red) and base nitrogen (blue) of A72 and C69.

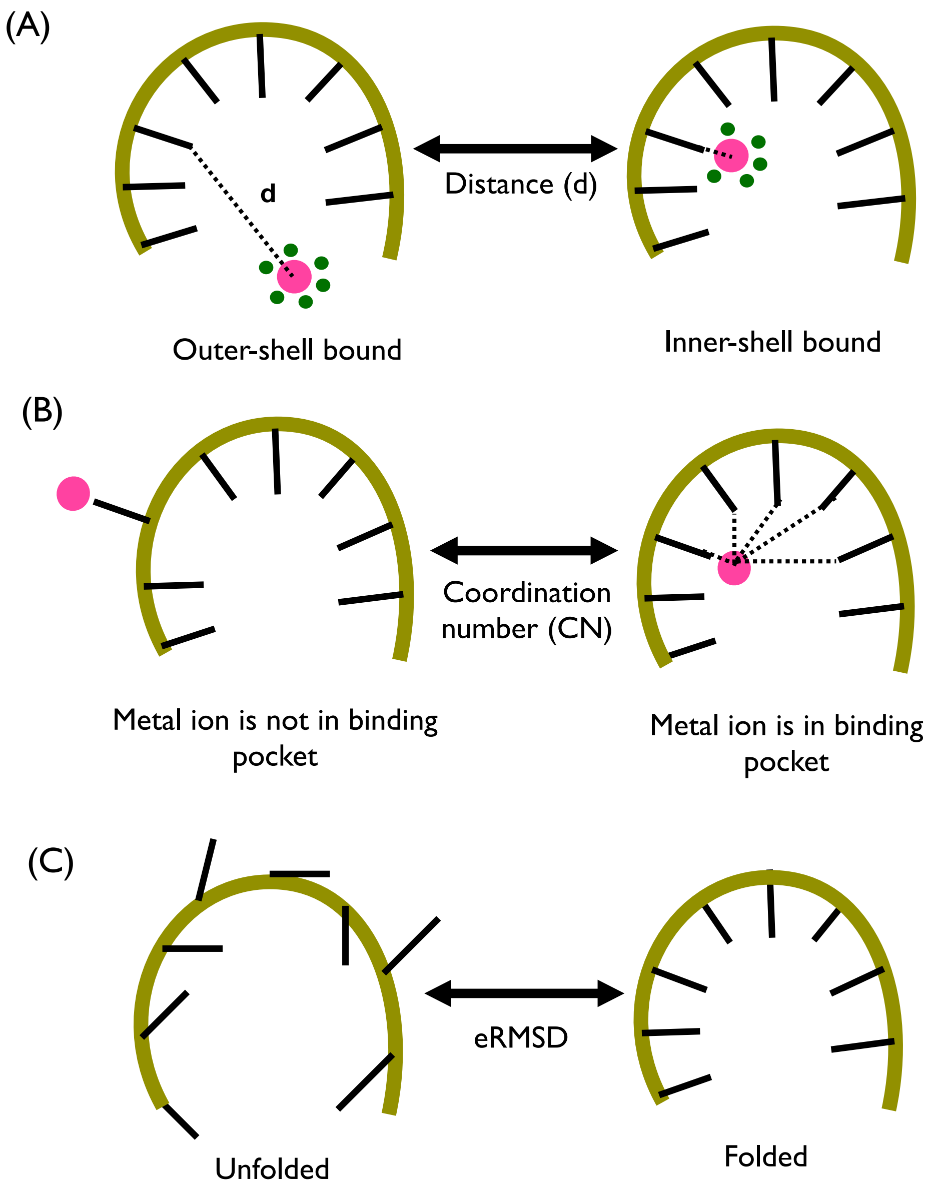


Figure S2: Schematic representation of collective variables (A) $d$, (B) $CN$, and (C) $eRMSD$ used in the WT-MetaD simulations to construct the folding FES of the L4 tetraloop. The backbone and nucleobases are shown in tan and black color. Magenta and green beads represent the metal ion and water molecules, respectively. The broken black lines show the contacts between the metal ion and the RNA atoms.

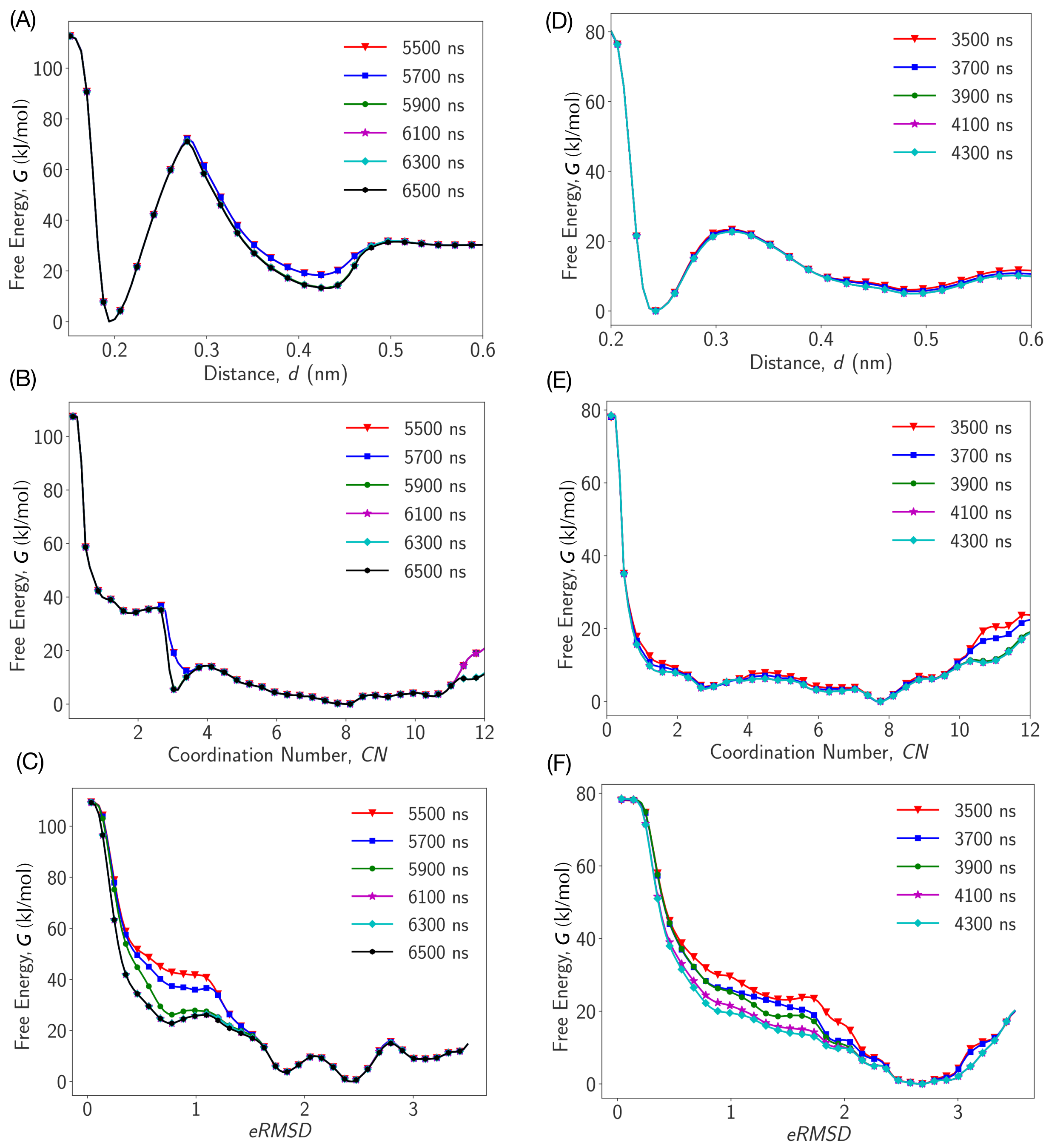


Figure S3: Convergence for WT-MetaD simulations: Free energy surface (FES), $G$ as a function of (A) distance, $d$, (B) coordination number, $CN$ and (C) $eRMSD$ for $Mg^{2+}$. Same for $Ca^{2+}$ is shown in (D), (E) and (F). No noticeable change is observed in the FES during the final $\approx$ 1000 (800) ns of the simulations for $Mg^{2+}$ ($Ca^{2+}$). The annotations represent the length of the simulation data used for computing FES.

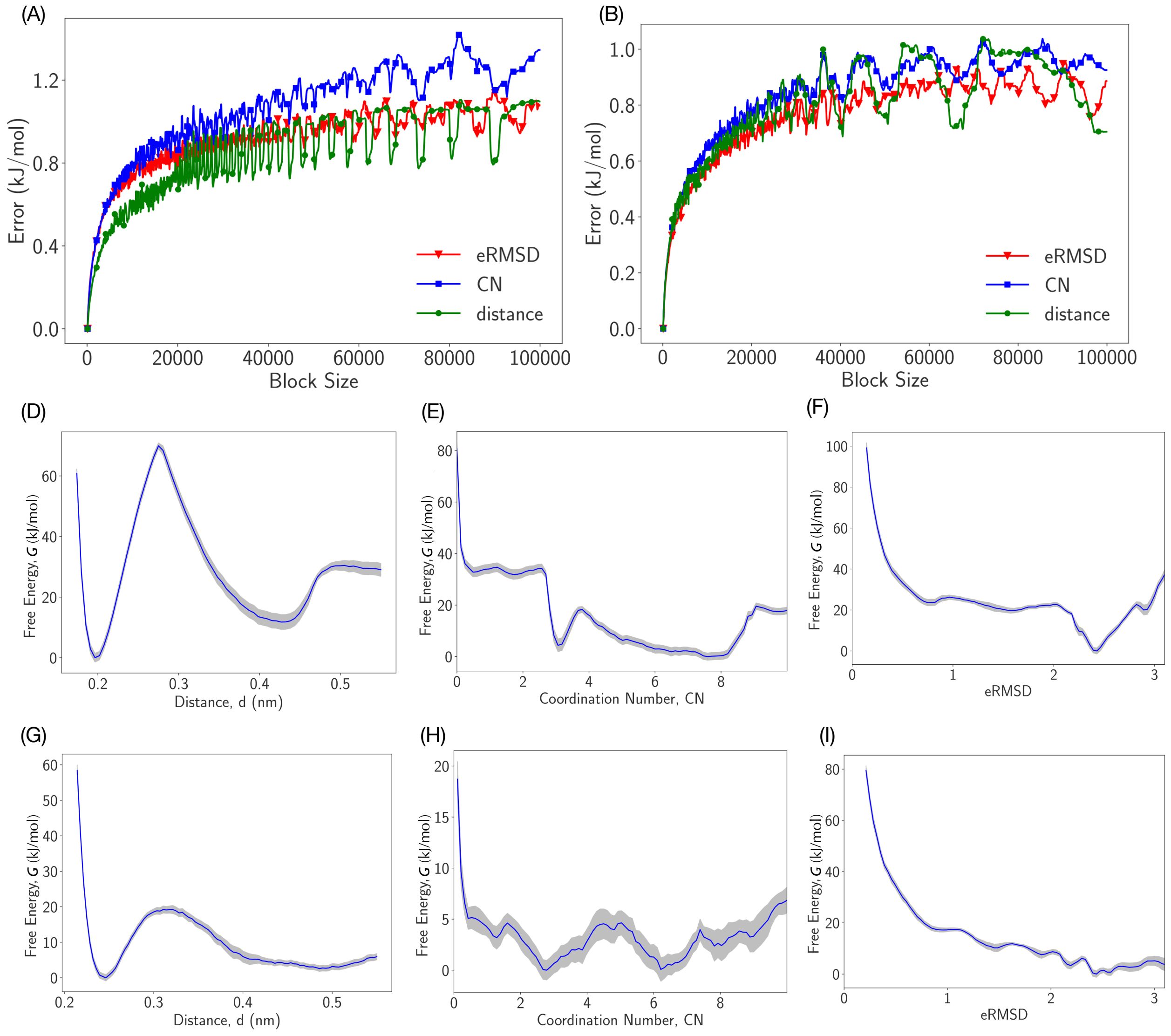


Figure S4: Error bars for three CVs ($d1$, $CN$ and $eRMSD$) are measured using block analysis for (A) $Mg^{2+}$-Tetraloop system, (B) $Ca^{2+}$-Tetraloop system. Errors in free energy profiles of above mentioned CVs are represented in the silver filled region and the average free energy profiles are shown in blue curve for (D,E,F) $Mg^{2+}$-Tetraloop system, (G,H,I) $Ca^{2+}$-Tetraloop system.

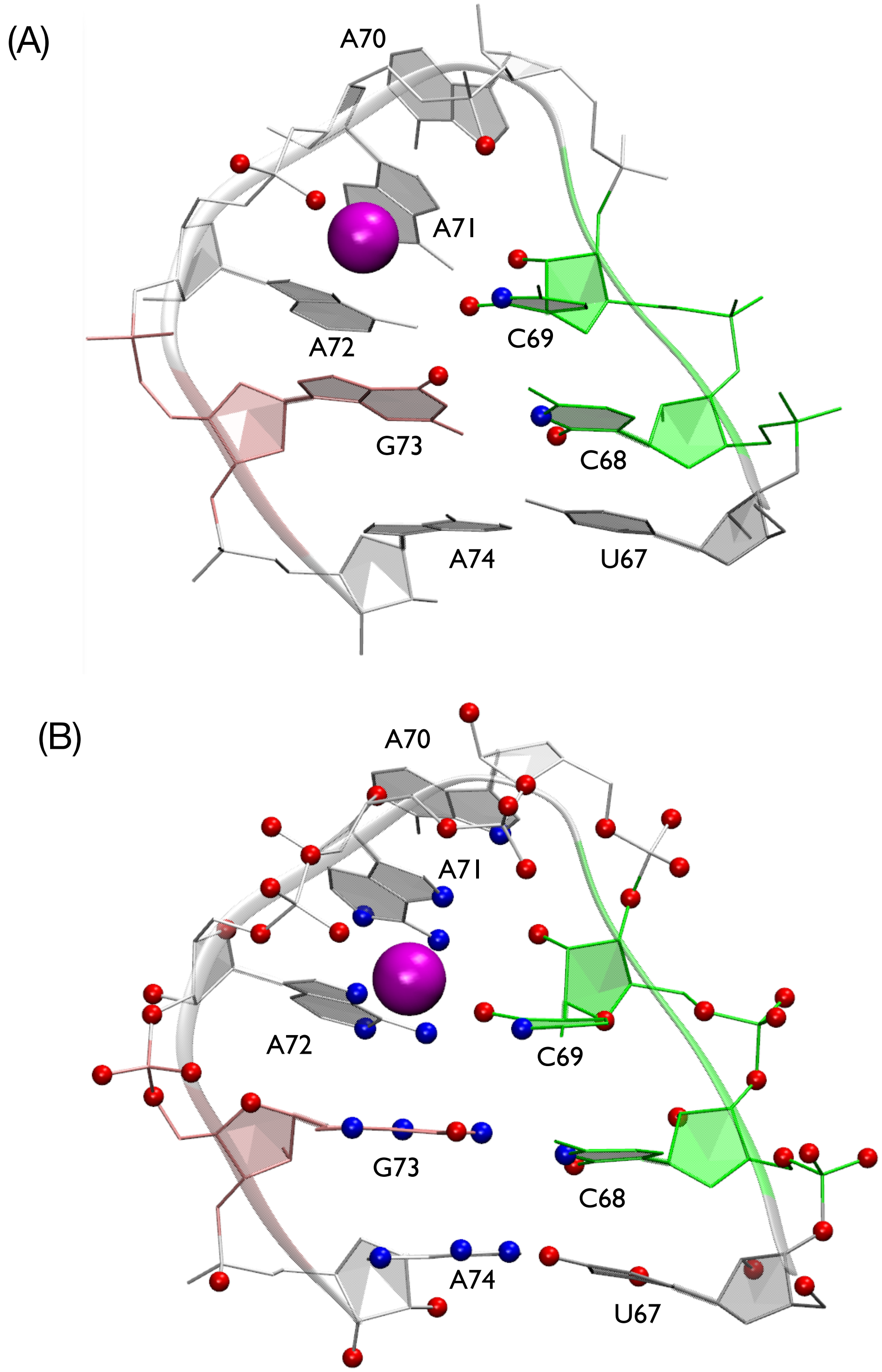


Figure S5: The heavy atoms of tetraloop that are involved in inner-shell coordination with metal ions, (A) $Mg^{2+}$ (pink sphere) and (B) $Ca^{2+}$ (pink sphere) are shown with red (oxygen) and blue (nitrogen) spheres. The L4 tetraloop is shown in the NewRibbon and CPK representation. $Mg^{2+}$ ($Ca^{2+}$) forms inner-shell contacts with 9 (58) RNA atoms.

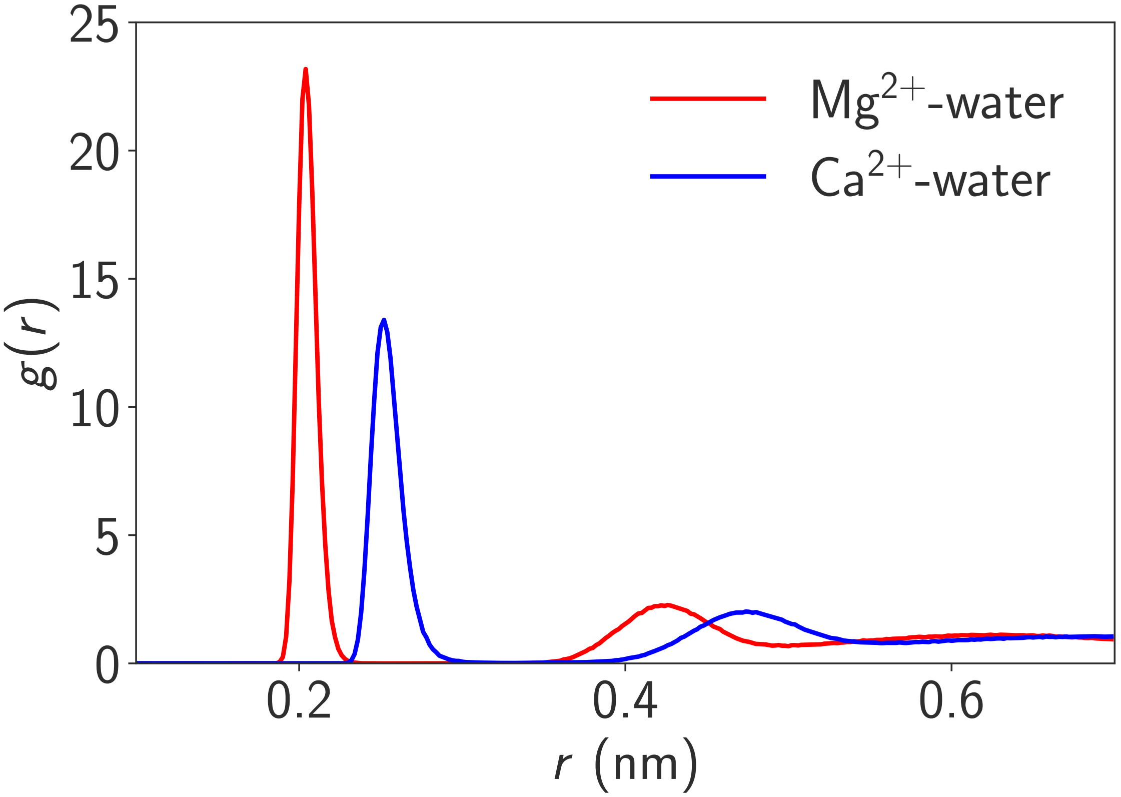


Figure S6: Radial distribution function between $M^{2+}$ ion (M = Mg, Ca) and oxygen atoms of water molecules in water-ion system.

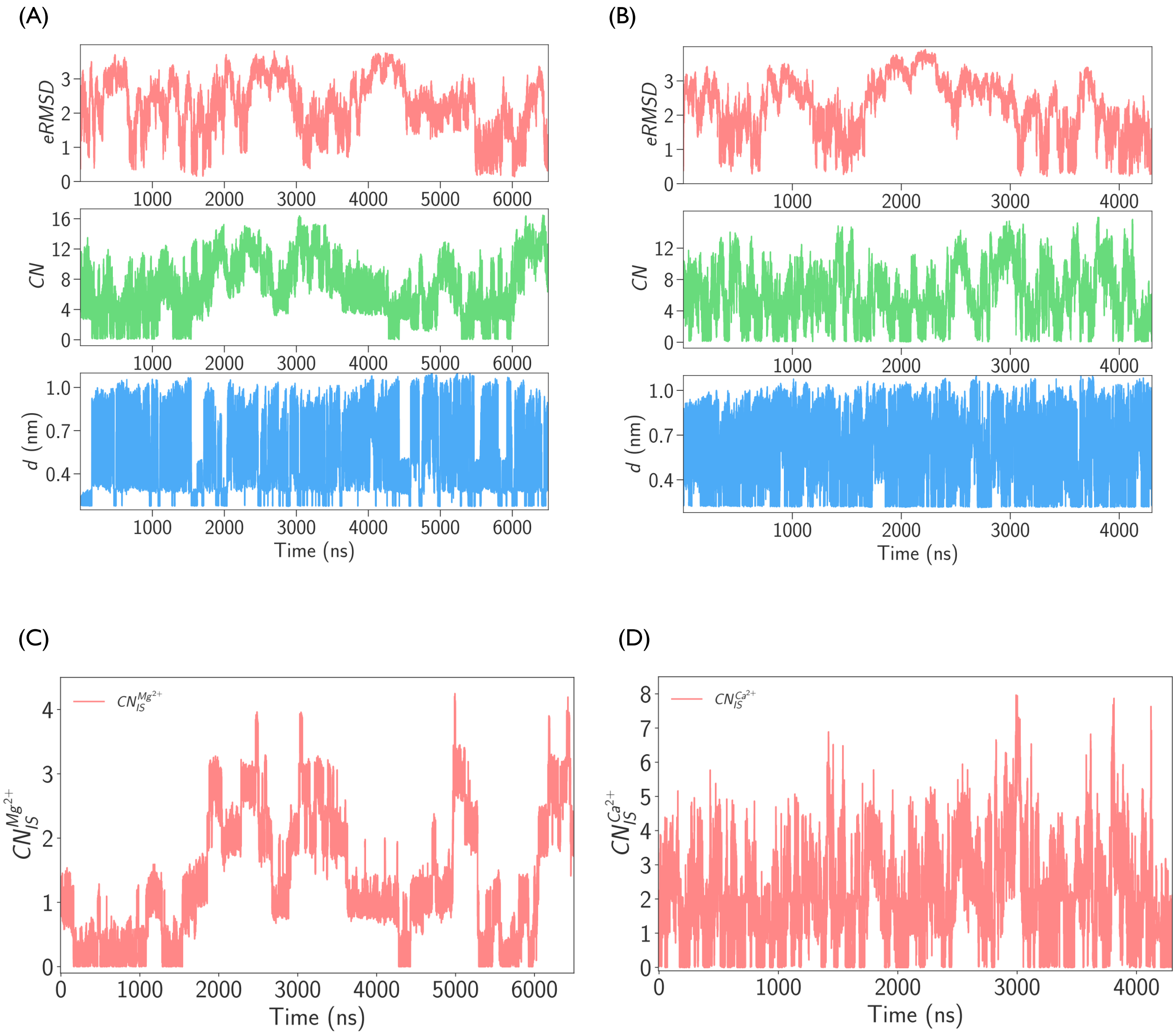


Figure S7: Collective variables, $eRMSD$, $d$ and $CN$ as a function time in WT metadynamics simulation for (A) $Mg^{2+}$, (B) $Ca^{2+}$. Here, $eRMSD \leq 1$ ($eRMSD > 1$) corresponds to the native-like folded (unfolded) state of the L4 tetraloop. If $d \leq 0.3$ nm ($d > 0.3$ nm), the metal ion is inner-shell (outer-shell) coordinated with atom A72-O2P. $CN$ determines the number of contacts around the metal ion. Coordination number between $M^{2+}$ ion and heavy atoms (oxygen and nitrogen) of tetraloop present in the first solvation shell of $M^{2+}$ ion, $CN_{IS}^{M^{2+}}$ (M = Mg, Ca) as a function of time for (C) $Mg^{2+}$, (D) $Ca^{2+}$.

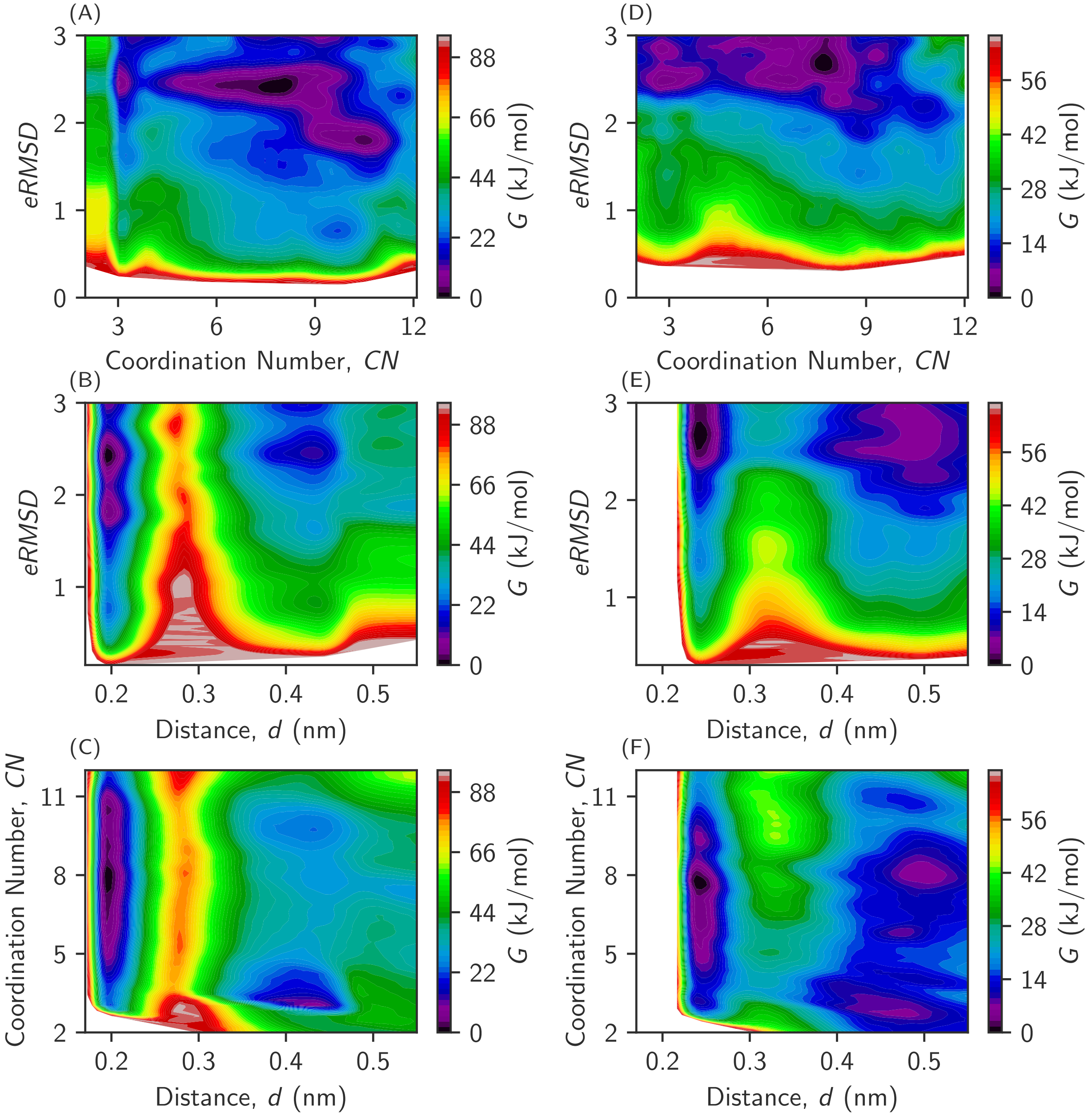


Figure S8: (A) $Mg^{2+}$-tetraloop FES as a function of (A) $eRMSD$ and $CN$, (B) $eRMSD$ and $d$, and (C) $CN$ and $d$ for $Mg^{2+}$. Similar FES for $Ca^{2+}$-tetraloop system are shown in (D), (E) and (F). The color bar represents the free energy, $\Delta G$.

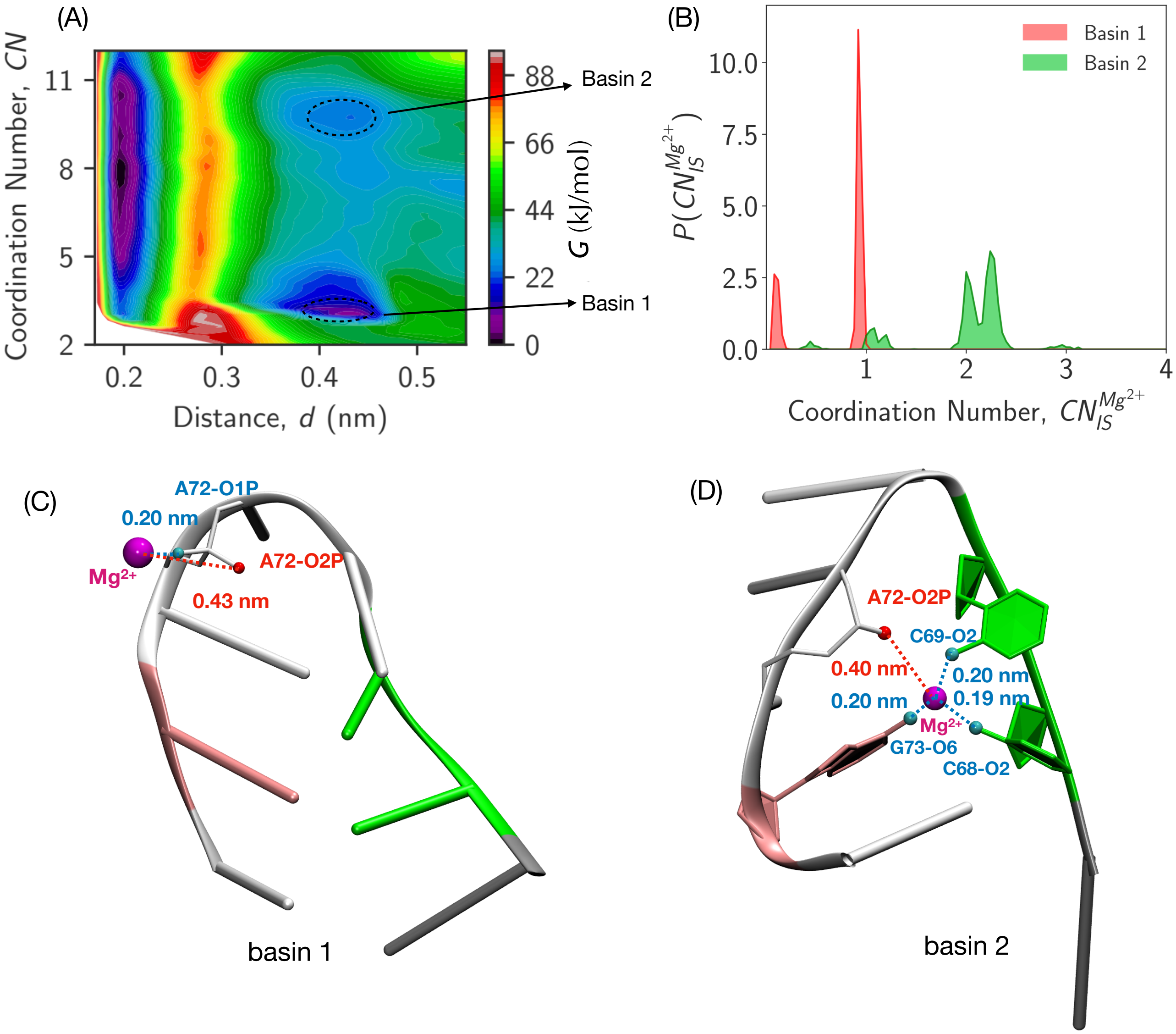


Figure S9: (A) FES as a function of $CN$ and $d$ for $Mg^{2+}$ where basin 1 ($CN \approx 3$) and 2 ($CN \approx 10$) are the most populated outer-shell coordinated structures. (B) Frequency of inner-shell contact formation with RNA atoms around $Mg^{2+}$ ion, $P(CN_{IS}^{Mg^{2+}})$ for basin 1 and 2 structures. Representative structures from basin 1 and 2 are shown in (C) and (D). Red broken line indicates the OS coordination distance ($\approx 0.4$ nm) between $Mg^{2+}$ (magenta beads) and A72-O2P atom (red sphere). Blue broken lines describe the IS coordination distance (0.2 nm) between $Mg^{2+}$ (magenta beads) and other IS-coordinated atoms A72-O1P, C69-O2, C68-O2, G73-O6 atom etc (green spheres). Basin 1 (Basin 2) structures form one (three) IS contact(s). Tetraloop conformations are givem in NewCartoon, Licorice and NewRibbons representations.

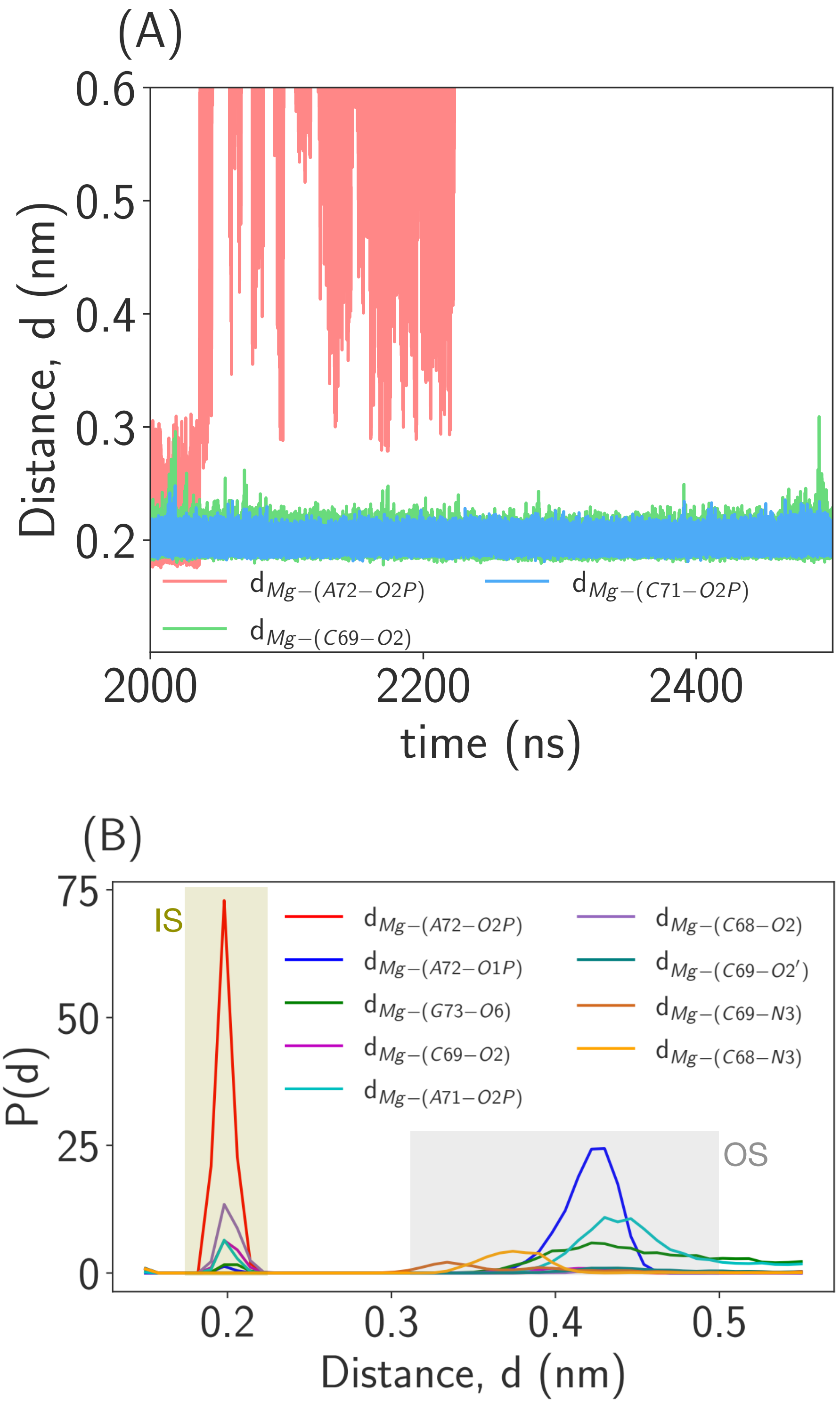


Figure S10: (A) Distance between $Mg^{2+}$ and RNA atoms, $d$ is shown as a function of time. $d_{Mg^{2+}-(A72-O2P)}$ (pink), $d_{Mg^{2+}-(A71-O2P)}$ (cyan) and $d_{Mg^{2+}-(A69-O2)}$ (green) are the distance between $Mg^{2+}$ and RNA atoms A72-O2P, A71-O2P and A69-O2, respectively. (B) Probability of inner-shell (IS) contact formation by each IS-coordinated atom for $Mg^{2+}$-tetraloop system.

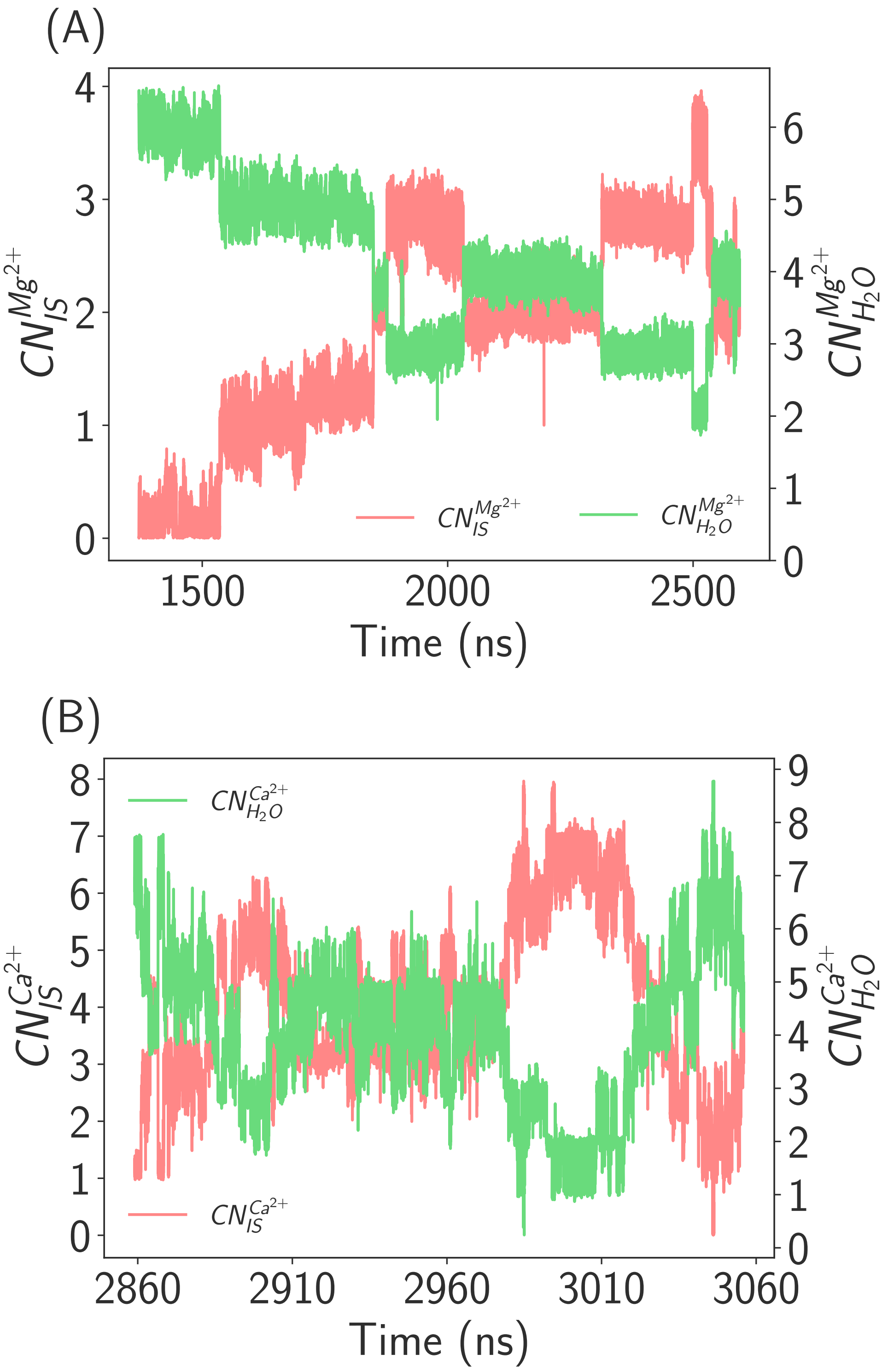


Figure S11: Representative simulation trajectories showing the removal of water from the first solvation shell of the ion and formation of IS contacts with the RNA atoms as a function of time for (A) $Mg^{2+}$ and (B) $Ca^{2+}$. $CN_{IS}^{M^{2+}}$ and $CN_{H_2O}^{M^{2+}}$ are the number of RNA atoms and water around metal ions, $M^{2+}$ ($Mg^{2+}$ and $Ca^{2+}$) in the first solvation shell.

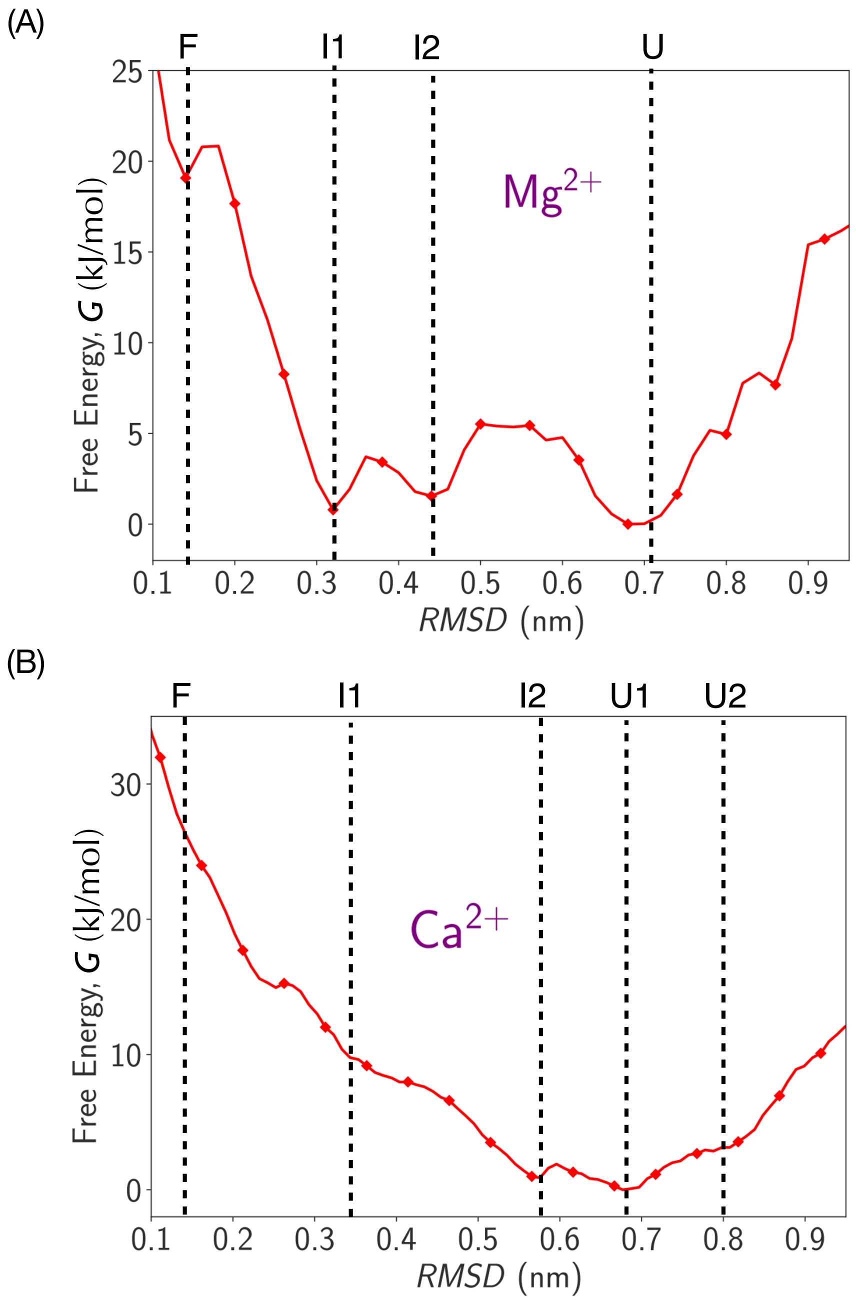


Figure S12: Free energy surface (FES) projected on RMSD for L4 tetraloop in presence of (A) $Mg^{2+}$ and (B) $Ca^{2+}$.

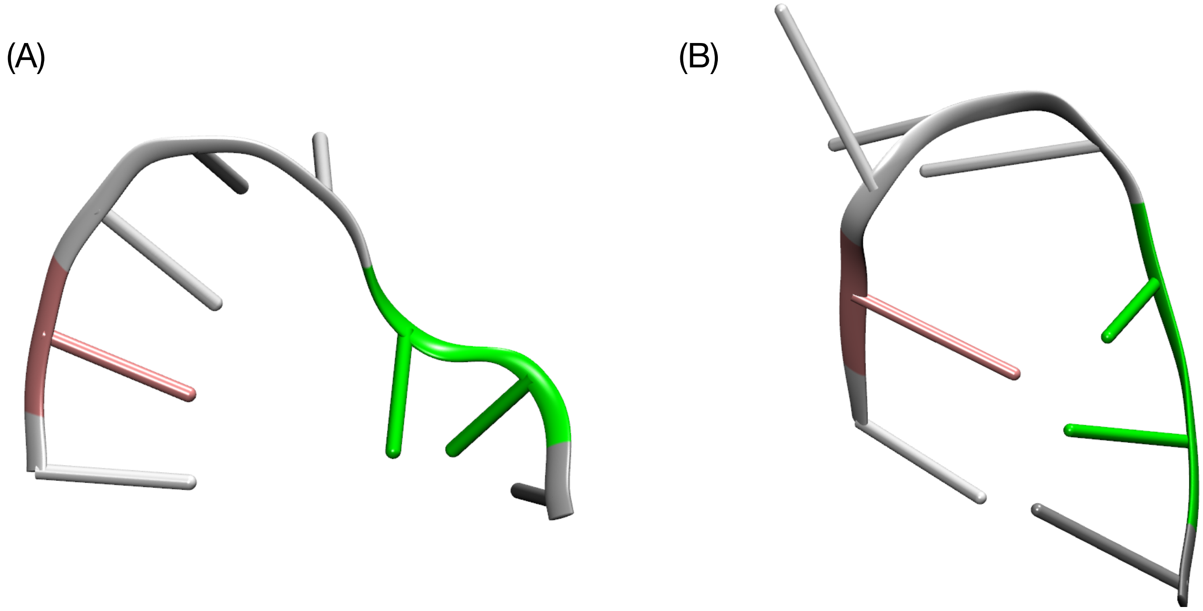


Figure S13: L4 tetraloop structures similar to the experimentally observed structures (A) E-type structure and (B) S-type structure found in I1 and I2 states.

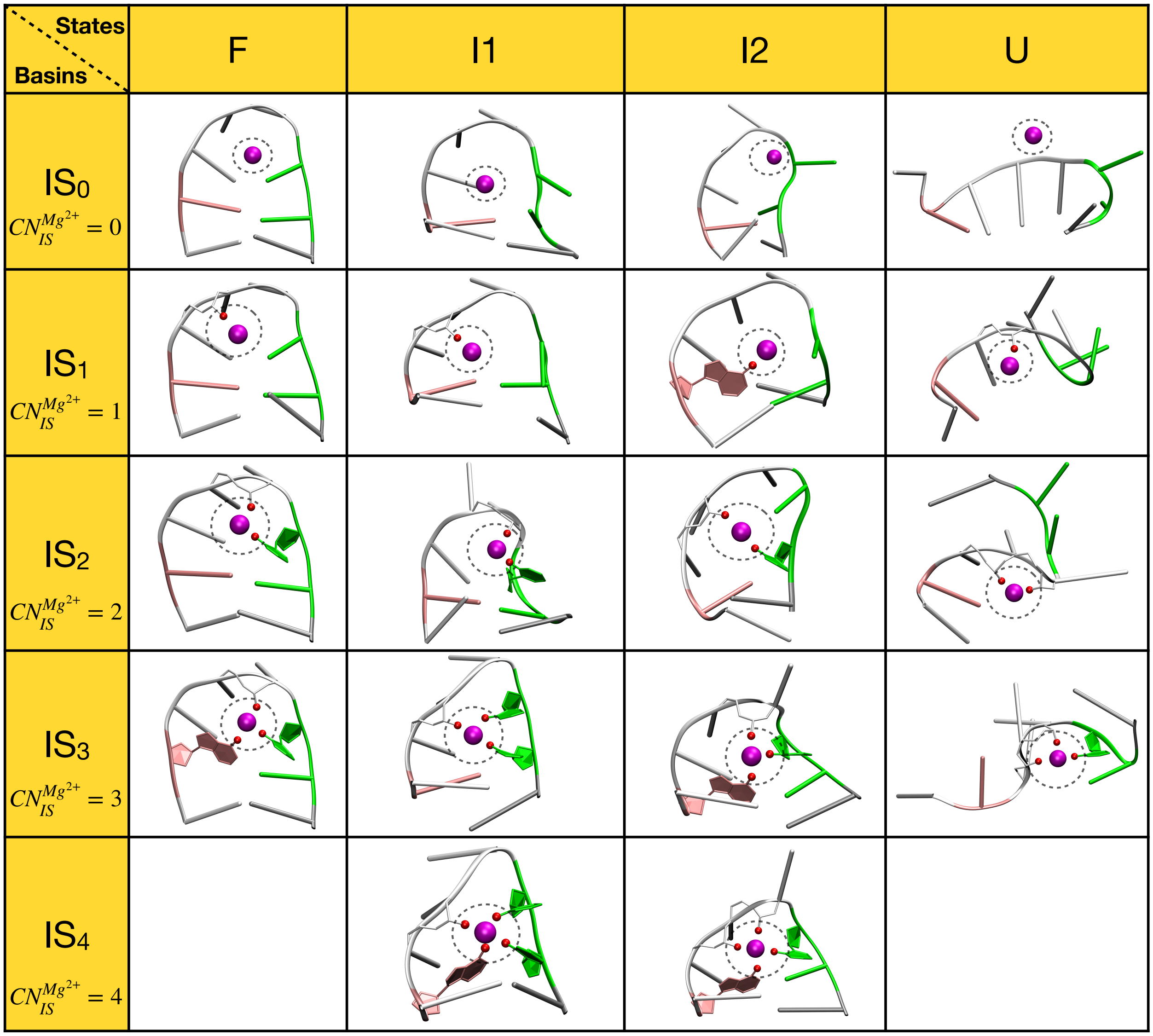


Figure S14: Classification of different states based upon RMSD and number of inner-shell contacts formed for $Mg^{2+}$. Structures $IS_0$ to $IS_4$ have 0 to 4 inner-shell contacts between $Mg^{2+}$ ion and electronegative atoms of RNA residues. IS-contact formation between $Mg^{2+}$ ion (magenta sphere) and RNA atoms (red spheres) is highlighted with the black circle.

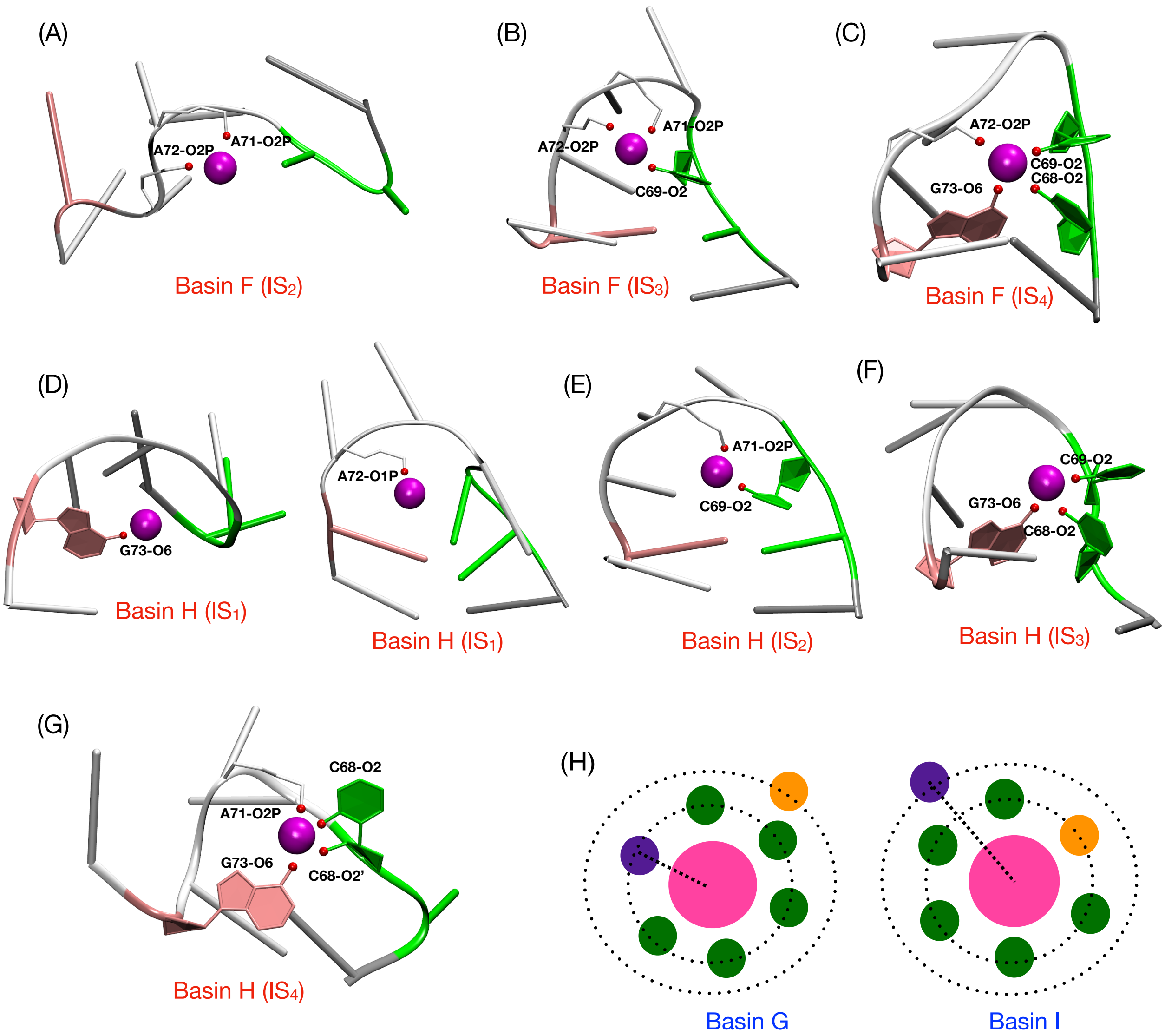


Figure S15: Respresentative non native IS coordinated basin F structures from simulation of (A) $IS_2$ conformation, (B) $IS_3$ conformation, (C) $IS_4$ conformation. Respresentative non native IS coordinated basin H structures from simulation (D) two different $IS_1$ conformations, (E) $IS_2$ conformation, (F) $IS_3$ conformation, (G) $IS_4$ conformation. (H) Schematic representations of the structures from the basin G and I are shown here. The water molecule in yellow (purple) is in the second (first) solvation shell in basin G. After the exchange process, the water molecule in yellow (purple) is in the first (second) solvation shell in basin I.

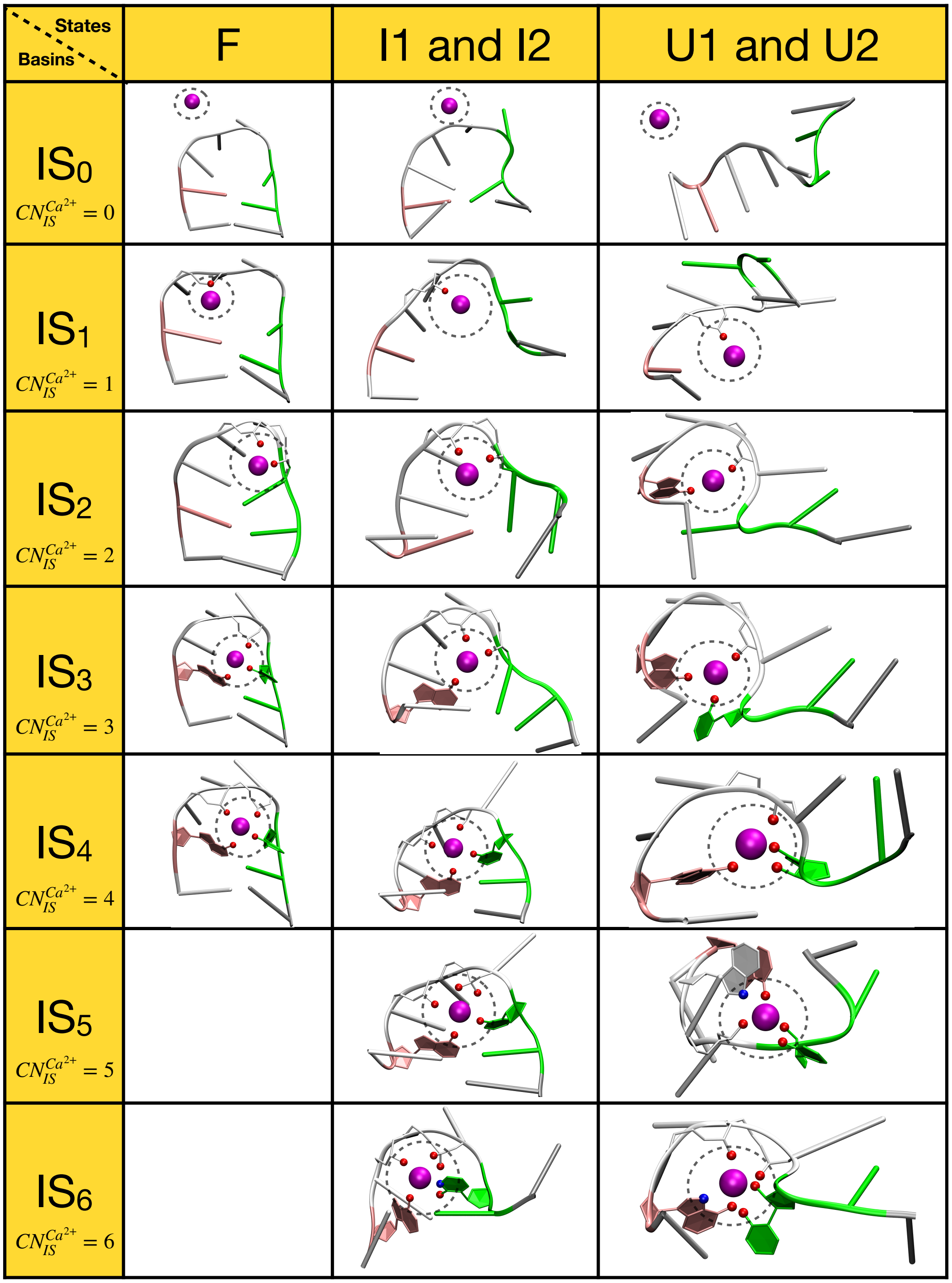


Figure S16: Classification of different states based upon RMSD and number of inner-shell contacts formed for $Ca^{2+}$. Structures $IS_0$ to $IS_6$ have 0 to 6 inner-shell contacts between $Ca^{2+}$ ion and electronegative atoms of RNA residues. IS-contact formation between $Ca^{2+}$ ion (magenta sphere) and RNA oxygen atoms (red spheres) and RNA nitrogen atoms (blue spheres) are highlighted with the black circle.

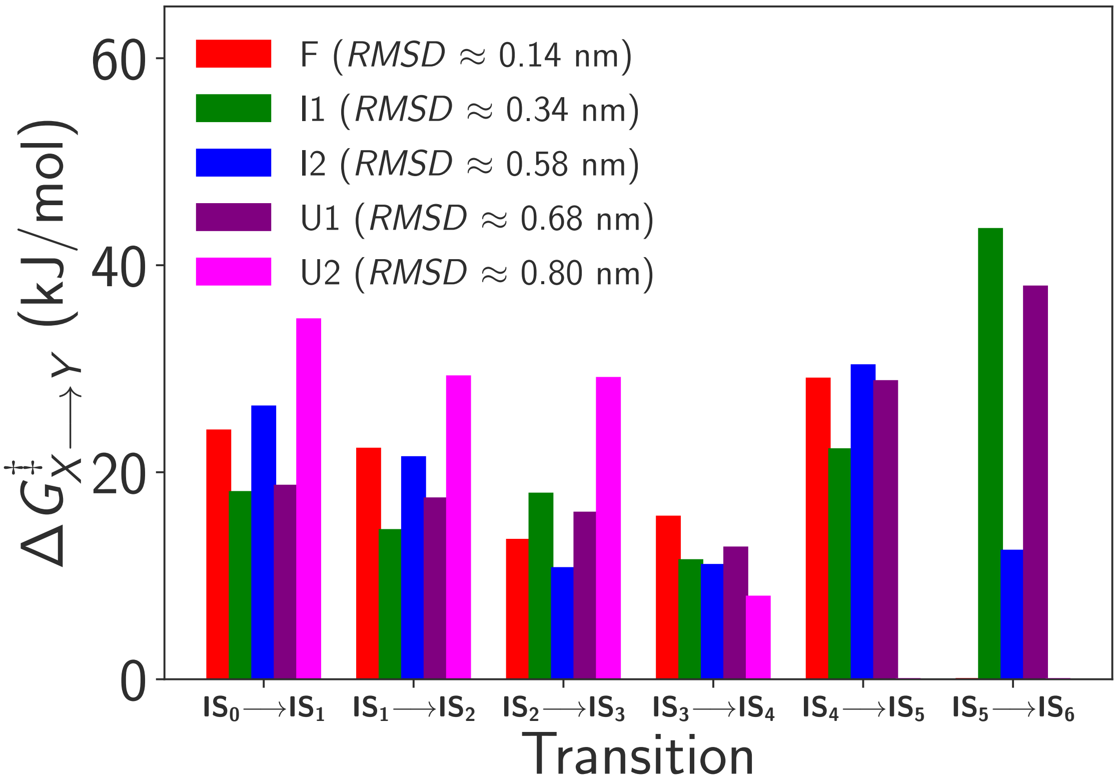


Figure S17: The free energy barrier, $\Delta G^{\ddagger}_{X \to Y}$ associated with the transition from basin $X$ (= $IS_0$, $IS_1$, $IS_2$, $IS_3$, $IS_4$, $IS_5$) to $Y$ (= $IS_1$, $IS_2$, $IS_3$, $IS_4$, $IS_5$, $IS_6$) for F (red), I1 (green), I2 (blue), U1 (purple) and U2 (magenta) state for $Ca^{2+}$-tetraloop system.

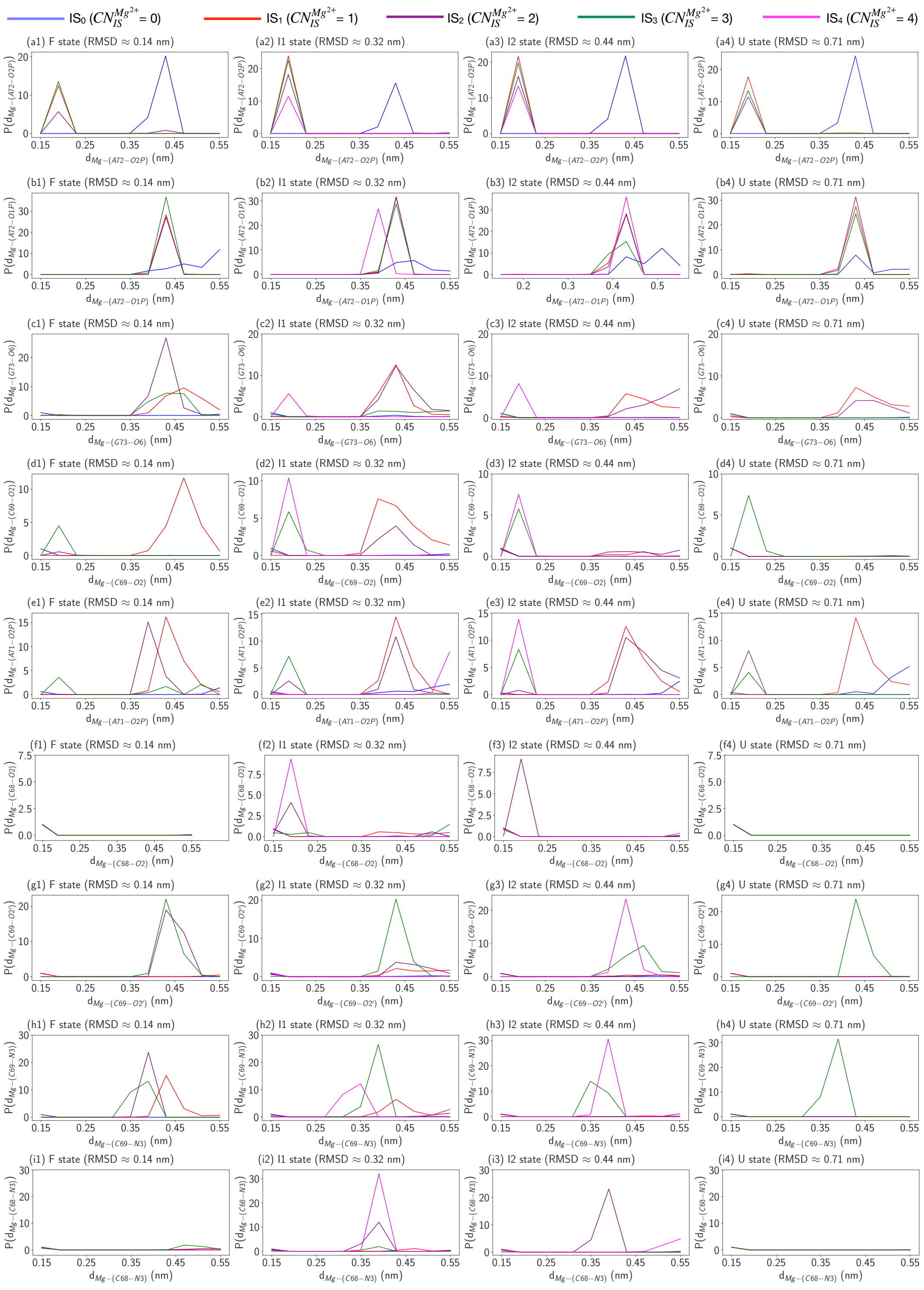


Figure S18: Probability of IS contact formation (at d ≈ 0.2 nm) by 9 IS-coordinated atoms (a-i) in $IS_0$ (blue), $IS_1$ (red), $IS_2$ (purple), $IS_3$ (green) and $IS_4$ (magenta) basins for F, I1, I2 and U states for $Mg^{2+}$-tetraloop system.

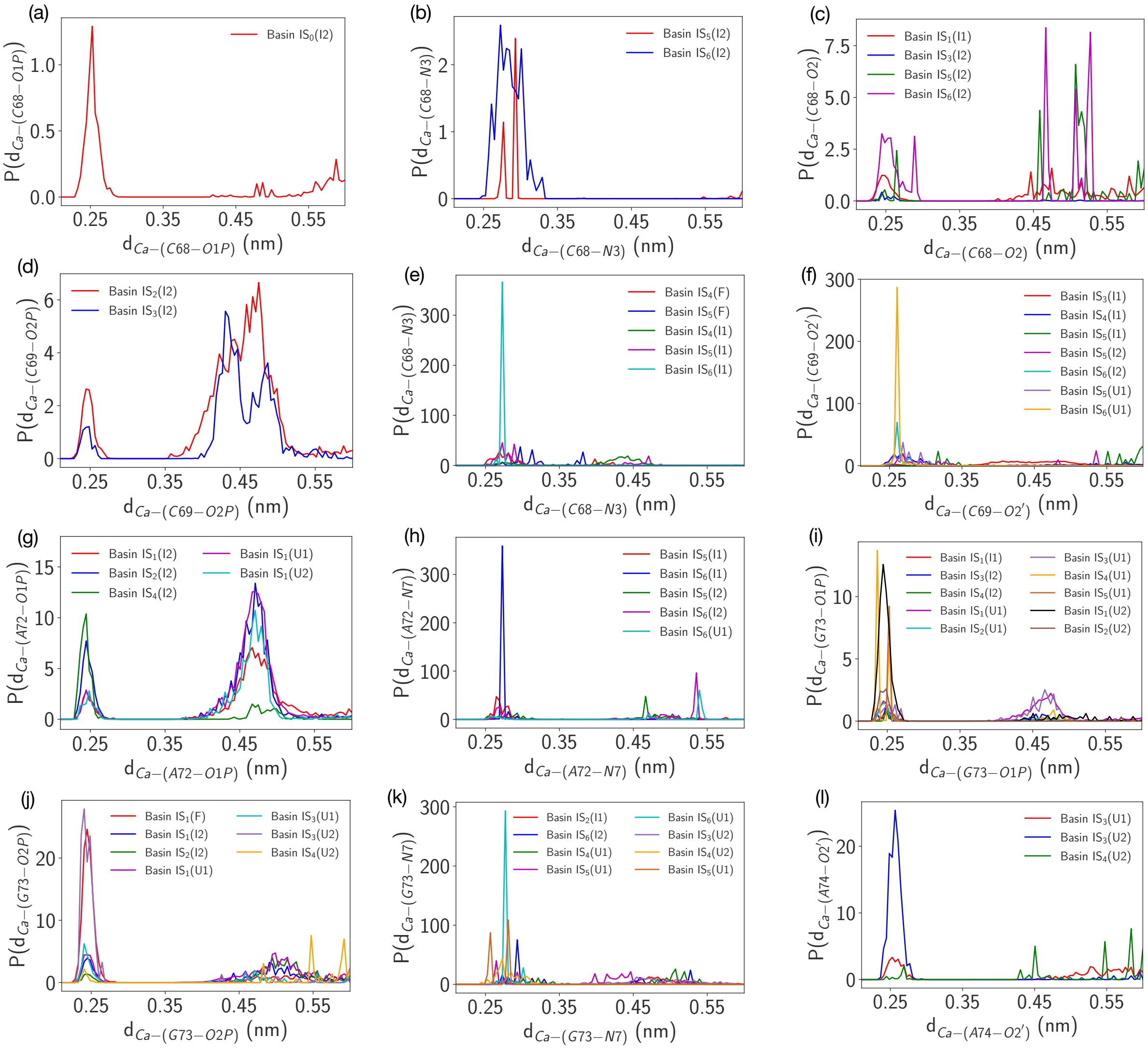


Figure S19: Probability of IS contact formation (at d $\approx$ 0.25 nm) by 12 IS-coordinated atoms (a-l) in different basins marked in legends for $Ca^{2+}$-tetraloop system.

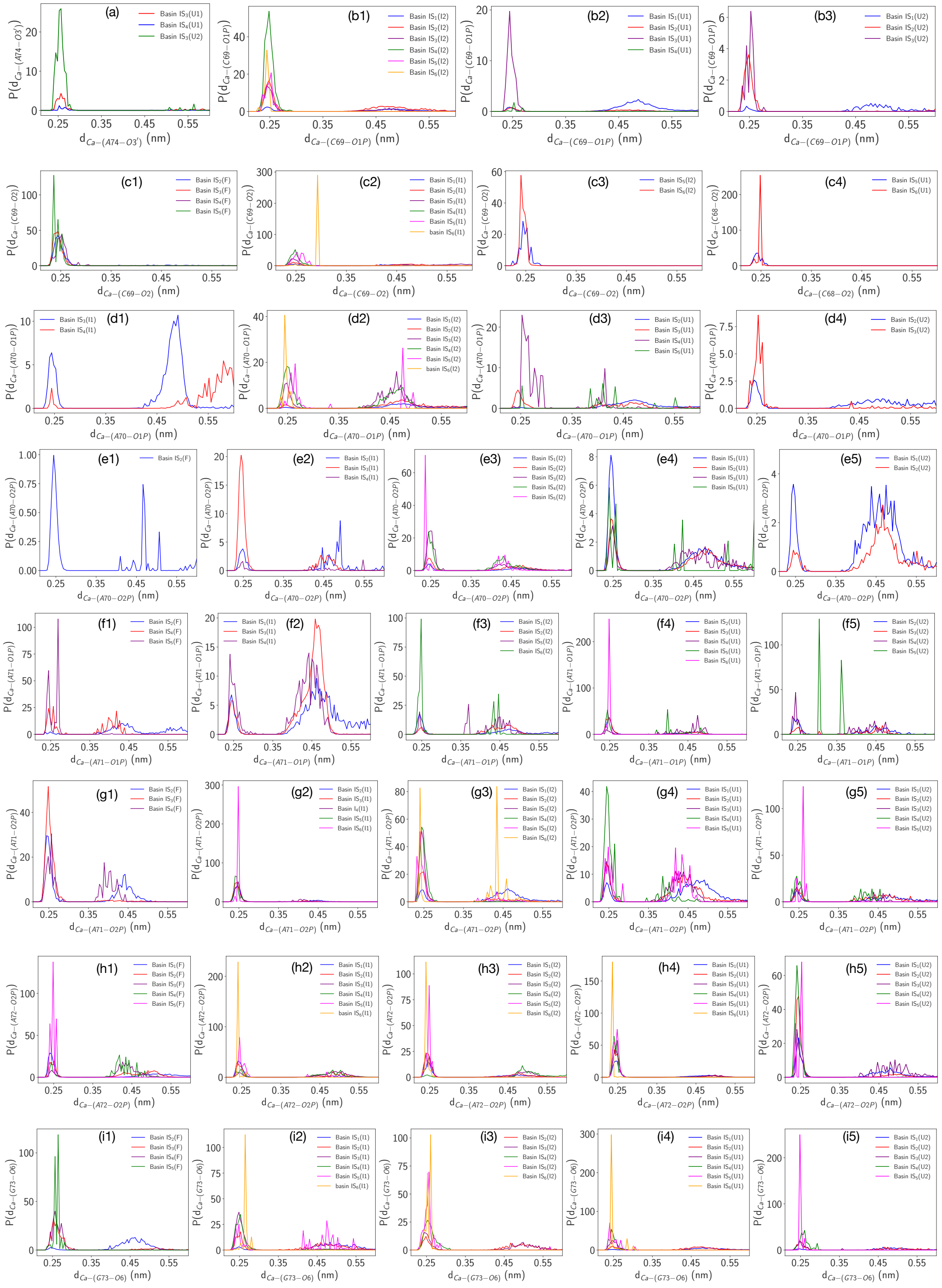

Figure S20: Probability of IS contact formation (at d ≈ 0.25 nm) by 9 IS-coordinated atoms (a-i) in different basins marked in legends for $Ca^{2+}$-tetraloop system.
